# ProteinFlux: accurate, rapid and scalable generative prediction of protein dynamics driven by post-translational modifications

**DOI:** 10.64898/2026.05.06.721822

**Authors:** Qiuting Qian, Jiahua Peng, Dongge Ma, Kaining Liu, Yihan Cheng, Yanqi Deng, Jingjing Zhao, Shen Su, Yifei Yao, Yuan Qu, Rao Fu, Jinliang Liu, Make Zhao, Yueming Xiao, Keying Wang, Yizhen Wu, Yijun Wang, Qiufan Xu, Jiarui Wang, David C. Hay, Yuehai Ke, Yong Wang, Michael J. Shipston, Ying Chi

## Abstract

The function of proteins, the building blocks of life, in health and disease depends not only on their 3D-conformational states but most importantly on the dynamic transition between states controlled by a wide array of post-translational modifications (PTMs). Recent major advances have been made in our ability to predict static 3D structures; however, understanding and predicting the impact of PTMs on protein conformational dynamics remains a major question and challenge in the field. Molecular dynamics (MD) simulation remains the major computational approach for studying protein dynamics. However, the high computational cost, lack of integration of PTMs as conditioning inputs and inefficient generation of continuous protein dynamics largely precludes PTM-regulated conformational dynamics and the study of slow conformational processes. To address this critical bottleneck, we developed ProteinFlux, a flow-matching generative framework that links PTM-conditioned conformational dynamics to evolutionary constraints encoded by PTM sites. Evolutionary information plays a critical role in capturing conformational dynamics beyond sequence identity, and PTM sites inherently encode evolutionary constraints critical to protein functional regulation. We therefore built FluxSite, a dual-modal PTM site predictor that integrates sequence evolutionary information and 3D structural features to generate a continuous conditional signal encoding conservation and functional importance for each predicted site. FluxSite achieves robust generalization across 18 PTM types and 30 disease-associated proteomes. ProteinFlux generates phosphorylation-conditioned, all-atom conformational trajectories across diverse protein fold classes, faithfully reproducing both thermodynamic properties such as free energy landscapes and kinetic features such as conformational transition pathways. It outperforms state-of-the-art predictors while achieving inference speeds several orders of magnitude faster than traditional MD. In addition, we introduce DynaMo-phos, a benchmark dataset of phosphorylated protein MD simulations. Together, ProteinFlux, FluxSite and DynaMo-phos provide a scalable, high-throughput platform for elucidating PTM-driven conformational mechanisms, with potential applications across allosteric drug design, functional annotation of disease-associated modifications and mechanism-guided therapeutic development.

## 1 Introduction

Proteins are the primary effectors of biological function, mediating processes ranging from enzymatic catalysis and signal transduction to molecular transport and structural support. Their functional diversity arises not from static three-dimensional structures alone, but from the conformational ensembles they populate and the temporal order of transitions between states ^1^. Disruption of these dynamic processes ^2,3^ is increasingly recognized as a driver of disease, positioning protein dynamics as both a fundamental biophysical property and a therapeutic target^4^. Understanding and predicting the relationship between protein sequence, post-translational modification (PTM) state, and conformational dynamics is therefore a central challenge in structural biology. Deep learning methods, led by AlphaFold2^5^ and related approaches ^6–10^, have transformed protein structural biology, achieving near-experimental accuracy^11,12^ for structure prediction and extending to multi-chain complexes, nucleic acids, small molecules, and covalent modifications. However, these advances address only the static dimension of this challenge. A critical gap remains in methods for predicting how protein conformational landscapes shift in response to specific biochemical perturbations, particularly those introduced by PTMs^13–16^. Furthermore, current methods struggle to generate the temporally ordered trajectories that capture how these perturbations propagate through the protein structure over time.

Molecular dynamics (MD) simulation remains the principal computational approach for studying protein dynamics. By modelling inter-atomic interactions with physics-based force fields, MD provides conformational trajectories spanning femtosecond-to-millisecond timescales and has been widely applied to protein folding, ligand binding, and allosteric mechanisms^17–19^. However, its substantial computational cost precludes genome-scale analyses or the characterisation of slow biological processes^20–22^. Recent deep generative models^23–28^ have begun to address this efficiency bottleneck by learning conformational sampling directly from data. Among these, BioEmu^25^ generates Boltzmann-distributed conformational ensembles with notable accuracy but produces independent samples without temporal ordering and does not support conditioning on biochemical perturbations such as PTMs. These methods, however, suffer from two fundamental limitations. Most methods generate only a set of disconnected structures to form a conformational ensemble, lacking explicit temporal ordering and physical time evolution. As a result, critical kinetic information related to transition pathways, time scales, and dynamic relaxation processes is inevitably lost. More critically, none incorporates biophysical perturbations such as PTMs as explicit conditioning inputs, meaning that a model trained on unmodified trajectories will generate an ensemble regardless of the modification state of the protein. Furthermore, the timescale gap between microsecond-accessible MD simulations and the millisecond-and-beyond processes that govern protein function makes adequate conformational sampling prohibitively expensive. Traditional enhanced sampling techniques partially address the sampling problem but remain unable to efficiently generate the continuous dynamic processes needed for mechanistic interpretation ^20,21^. Together, these limitations highlight the absence of a unified framework capable of predicting the continuous conformational trajectories induced by specific post-translational modifications.

Modeling how localised chemical modifications reshape global protein dynamics presents two challenges. The model must maintain precise geometric awareness at the atomic level while learning how chemical modifications propagate across the entire protein scaffold. This propagation occurs through long-range intramolecular interactions and allosteric communication, ultimately altering conformational populations and reshaping the underlying free energy landscape. Equally limiting is the absence of suitable training data. Public MD datasets such as ATLAS^29^, MoDEL ^30^, and mdCATH^31^ provide extensive trajectory coverage for unmodified proteins but contain virtually no systematic representation of PTM-modified systems. Given that PTMs affect the majority of proteome proteins ^32–34^, this data gap represents a key bottleneck for developing PTM-aware generative models. A third challenge lies in identifying a biologically meaningful conditioning signal that captures the functional significance of PTM sites beyond their sequence position.

To address these challenges, we present ProteinFlux, a flow-matching generative framework^35–37^ that links PTM site prediction to PTM-conditioned conformational trajectory generation. We find that sequence identity alone is insufficient to distinguish the distinct dynamic distributions across protein datasets. Incorporating evolutionary information encoded by protein language models substantially improves the model’s ability to capture conformational dynamics across diverse datasets. This reveals a mutually reinforcing relationship between evolutionary information and data diversity. However, as general-purpose sequence language models, they implicitly compress broad evolutionary constraints without knowledge of which residues undergo specific post-translational modifications. This gap motivated the development of FluxSite, a dual-modal sequence-structure predictor that extracts PTM-specific evolutionary constraint signals from ESM representations and identifies PTM sites across 18 modification types. FluxSite produces discriminative functional embeddings that enable end-to-end prediction from sequence and structure inputs to PTM-induced conformational trajectories. FluxSite can also be applied independently to prioritise candidate modification sites for experimental validation. We validate the complete pipeline on the STREX variant of the BK potassium channel ^38–41^, an out-of-distribution system absent from all training data. ProteinFlux correctly identifies the experimentally validated regulatory phosphorylation site and generates conformational trajectories consistent with independent MD simulations. In addition, we introduce DynaMo-phos, a benchmark dataset of phosphorylated protein MD trajectories, to support training and standardised evaluation of PTM-aware generative models.

## 2 ProteinFlux architecture

ProteinFlux is a flow-matching generative framework for PTM-conditioned protein dynamics (Fig. 1). It produces temporally coherent, all-atom conformational trajectories that reflect the structural consequences of specific post-translational modifications. The base dynamics generator adopts an SE(3)-equivariant flow-matching architecture ^28^ and is pre-trained on large-scale MD simulations to learn generalizable principles of conformational change across diverse protein families (Fig. 1a). To incorporate PTM-specific information, a dedicated site prediction network, FluxSite, is trained to identify PTM sites from sequence and structural features (Fig. 1b). FluxSite integrates ESM2 sequence representations with ESM-IF structure representations to extract PTM-specific evolutionary constraint signals. Its learned latent representations serve as functional embeddings that encode site-specific biochemical context. These embeddings are injected into the dynamics generator via a lightweight PTM adapter (Fig. 1d), enabling ProteinFlux to learn a direct mapping from PTMs to their downstream conformational responses.

**Fig. 1.**
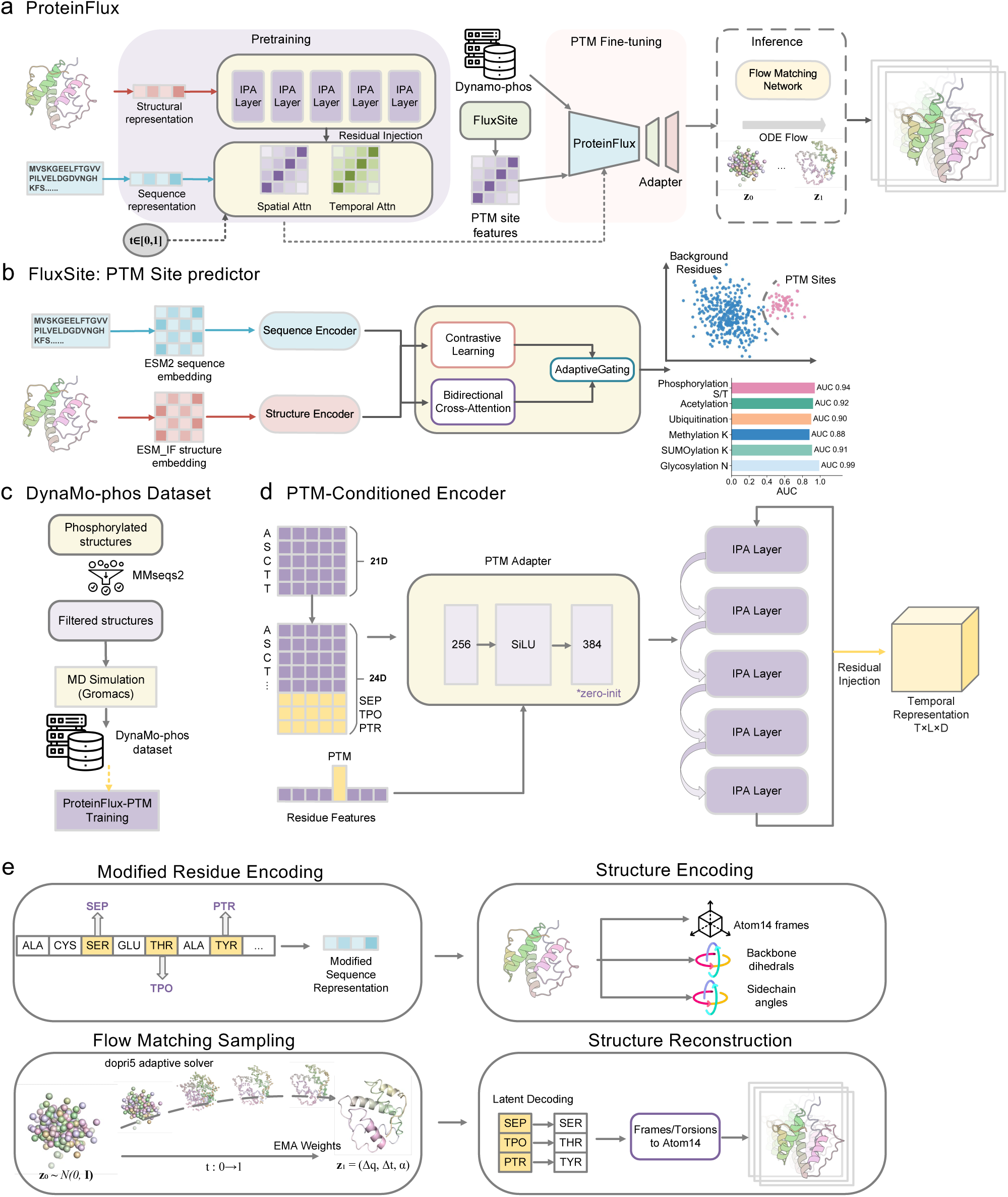
Overview of ProteinFlux architecture and workflow. (a) ProteinFlux framework. The model is pretrained with IPA layers and spatial-temporal attention modules. During PTM fine-tuning, FluxSite extracts PTM site features from DynaMo-phos and integrates them via a PTM adapter. At inference, the flow matching network maps a Gaussian prior (*z*_0_) to the target state (*z*_1_) through ODE-based sampling to generate PTM-conditioned conformational trajectories. (b) FluxSite: PTM site prediction module. ESM2 sequence encoder and ESM-IF structure encoder are fused via bidirectional cross-attention, with contrastive learning applied to learn discriminative site representations. (c) DynaMo-phos dataset. Phosphorylated protein structures are filtered via MMseqs2 sequence clustering ^42^, subjected to molecular dynamics (MD) simulations using GROMACS, and compiled into the training dataset for PTM-conditioned modeling. (d) PTM adapter. A zero-initialized adapter injects PTM-conditioned residue features as residual signals into each IPA layer without disturbing pre-trained weights. (e) Trajectory generation pipeline. Modified residues are encoded as distinct tokens: phosphoserine (SEP), phosphothreonine (TPO), and phosphotyrosine (PTR). The Dormand-Prince fifth-order (dopri5) adaptive ODE solver integrates the learned vector field from Gaussian prior (*z*_0_) to target latent state (*z*_1_) using exponential moving average (EMA)-weighted parameters. Latent representations are decoded through a frames/torsions-to-Atom14 reconstruction pipeline.

Existing MD databases predominantly contain unmodified proteins, leaving a critical gap for PTM-specific trajectory data. To address this, we constructed DynaMo-phos, a benchmark dataset focusing on phosphorylation, the most prevalent and functionally diverse PTM (Fig. 1c). The resulting dataset serves both as training data for ProteinFlux’s PTM conditioning module and as a standardized benchmark for evaluating PTM-driven dynamics prediction. Following pre-training, the model is fine-tuned on DynaMo-phos with PTM embeddings injected via the adapter module, learning to map modification-specific signals to conformational responses. To ensure generalization, training and test sets are split by sequence similarity (details in Methods). At inference, ProteinFlux takes a protein structure with annotated PTM sites and generates a temporally ordered trajectory of conformational states (Fig. 1e). Modified residues are explicitly encoded in the input representation, allowing the model to propagate PTM-specific effects throughout the generated dynamics.

## 3 Learning transferable protein dynamics with evolutionary representations

Protein function is governed not merely by the set of accessible conformational states, but critically by the kinetic pathways and chronological order of transitions between them. Regulatory mechanisms such as allosteric signal propagation^3^ and PTM-induced domain rearrangements inherently unfold through temporally ordered intermediates that dictate the ultimate functional outcome. While existing generative methods excel at sampling unordered structural ensembles, they systematically discard this vital temporal context. To build a robust computational foundation capable of recovering these continuous dynamics, we pre-trained ProteinFlux on an extensive molecular dynamics corpus encompassing 7,781 distinct protein systems ^29–31^ and over 13.3 ms of aggregated simulation time. We benchmarked the computational cost of ProteinFlux against AlphaFold3 and BioEmu by measuring GPU seconds required to generate an equivalent number of conformations. While AlphaFold3 predicts independent static structures and BioEmu samples unordered equilibrium states, ProteinFlux generates temporally ordered trajectories with richer dynamical content. Despite this, ProteinFlux achieves inference speeds one to two orders of magnitude faster than both methods across all tested sequence lengths. (Fig. 2e). ProteinFlux also exceeds BioEmu and MDGen on pairwise root mean square deviation (RMSD) correlation and per-target root mean square fluctuation (RMSF) correlation (Fig. 2f).

**Fig. 2.**
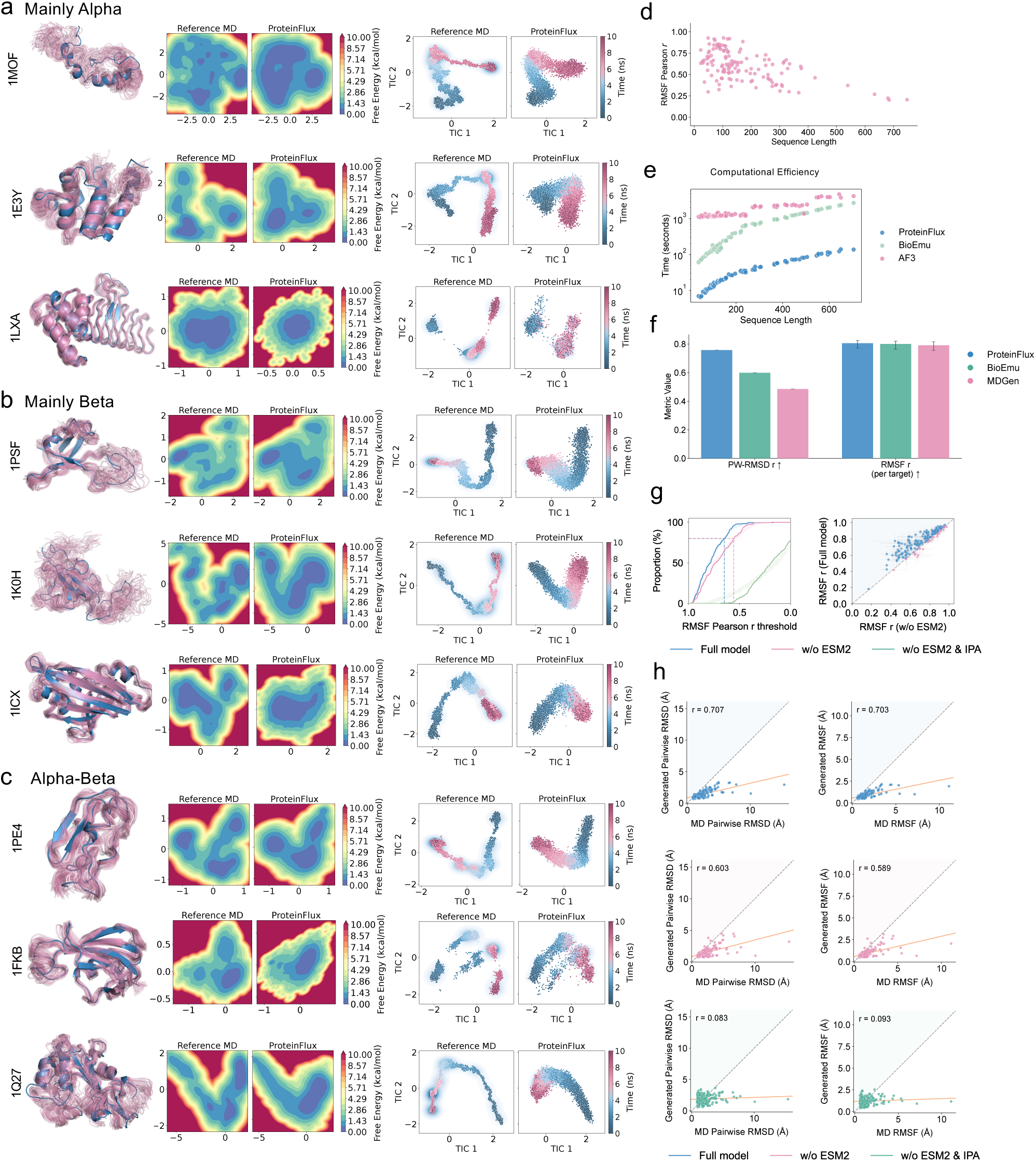
ProteinFlux generates temporally coherent protein dynamics. (a–c) Conformational analysis of the pre-trained ProteinFlux across protein structural classes. (a) Mainly alpha, (b) mainly beta, and (c) alpha-beta proteins, each illustrated with 3 representative proteins ordered by sequence length. The left panel shows superimposed reference molecular dynamics (MD) (blue) and ProteinFlux-generated (pink) conformational ensembles. The middle panels show free energy landscapes of the reference MD trajectory and ProteinFlux-generated trajectory, projected onto the first two principal components (PC1, PC2) of the reference MD trajectory. Free energy is in unit of kcal/mol. The right panels show time-lagged independent component analysis (tICA) projections onto the two slowest independent components (TIC1, TIC2) for the reference MD trajectory and ProteinFlux-generated trajectory, respectively, colored by simulation time (ns). (d) Per-protein RMSF Pearson correlation coefficient as a function of sequence length. (e) Computational efficiency comparison of ProteinFlux, BioEmu, and AlphaFold3 (AF3) for generating 250 conformational ensembles. Inference time is reported in seconds on a single GPU. (f) Benchmark comparison of ProteinFlux, BioEmu, and MDGen on pairwise C*_α_* RMSD correlation and per-target RMSF correlation. (g) Architectural ablation under multi-dataset joint training: full model, without ESM2 (w/o ESM2), and without both ESM2 and IPA (w/o ESM2 & IPA). Left: cumulative distribution of RMSF Pearson correlation. Right: per-protein RMSF correlation scatter plot. (h) Per-target scatter plots for three architectural variants (full model, w/o ESM2, and w/o ESM2 & IPA, from top to bottom). Left: pairwise C*_α_* RMSD correlation between generated and reference MD ensembles. Right: per-residue RMSF correlation.

We evaluated the model on the MoDEL test set using a strict 30% sequence identity cutoff to preclude data leakage. To ensure the generated ensembles are physically realistic, we assessed their thermodynamic equilibrium. We first reduced the conformational space via principal component analysis (PCA) and projected both reference and generated ensembles onto the first two principal components (PCs). Projections of the free energy landscapes onto the first two principal components demonstrate that ProteinFlux accurately samples the native conformational basins and equilibrium distributions across structurally diverse proteins, including representative systems from the Mainly Alpha (1MOF, 1E3Y, 1LXA), Mainly Beta (1PSF, 1K0H, 1ICX), and Alpha/Beta (1PE4, 1FKB, 1Q27) CATH classes (Fig. 2a–c). However, merely sampling thermodynamic states is insufficient for modeling true protein dynamics. To verify that ProteinFlux also captures the correct structural transitions over time, we applied time-lagged independent component analysis (tICA). The resulting projections onto the two slowest independent components confirm that the generated trajectories successfully recapitulate the kinetic transition pathways of the target systems(Fig. 2a–c). We further evaluated dynamical flexibility by comparing per-residue root mean square fluctuation (RMSF) profiles between reference and generated trajectories across diverse proteins in the MoDEL and ATLAS test sets (Extended Data Fig. 1).

Furthermore, we analyzed the sequence length boundaries imposed by the pre-training phase. Per-target root mean square fluctuation (RMSF) correlation remains strong for proteins under 300 residues but exhibits an expected decline for longer sequences (Fig. 2d). This is expected, as ProteinFlux was trained on cropped fragments of 256 residues, limiting its ability to capture long-range dynamical correlations in larger proteins. The effect of crop length on downstream performance is further analyzed in the PTM fine-tuning experiments (Fig. 5e). To ensure that the learned representations are invariant to global rotations and translations, ProteinFlux integrates the Invariant Point Attention (IPA) module^5,43^. This mechanism performs attention in local residue reference frames, providing an SE(3)-equivariant geometric inductive bias that enables the model to reason directly about spatial relationships between residues. Ablation experiments underscore the absolute necessity of this architecture. Removing the IPA module severely degrades performance, reducing pairwise root mean square deviation (RMSD) Pearson correlation and RMSF correlation(Fig. 2h).

Ablation of ESM2 from the jointly trained model led to marked performance degradation across both distributional and flexibility metrics (Fig. 2g,h), demonstrating that evolutionary representations are critical for multi-dataset dynamics generation. To dissect this effect, we examined the training strategy in detail. Jointly training on ATLAS and MoDEL at 100 ps resolution using amino acid type embeddings in place of ESM2 revealed negative transfer on ATLAS and MoDEL (Extended Data Fig. 2a), indicating that sequence-level representations alone cannot reconcile distributional differences between datasets with different simulation protocols. Replacing amino acid embeddings with ESM2 evolutionary representations eliminated this negative transfer, with joint training consistently matching or improving upon single-dataset performance (Extended Data Fig. 3a). Notably, this effect is specific to the multi-dataset setting. In single-dataset training, ESM2 provided limited or negligible improvement (Extended Data Fig. 3c), indicating that evolutionary representations are not universally beneficial but rather become essential when the model must reconcile heterogeneous data distributions. These findings establish that evolutionary representations are essential for learning transferable protein dynamics across heterogeneous datasets.

## 4 Identifying PTM sites across diverse modification types

While ESM2 evolutionary representations encode general sequence-structure-function constraints, they do not explicitly identify which residues undergo specific modifications or capture the distinct sequence contexts associated with different PTM types. To extract PTM-specific signals from evolutionary and structural information, we trained a dual-modal predictor, FluxSite, that fuses ESM2 sequence features with ESM-IF structural features to identify PTM sites and encode their biochemical identity into functional embeddings for conditioning the dynamics generator.

t-SNE visualization of the learned representations shows that ESM2 sequence features already achieve clear separation between PTM and background residues across representative PTM types (Fig. 3a), with the complete set of 18 types in Extended Data Fig. 4. However, fusing structural features consistently improves prediction performance beyond sequence alone (Fig. 3b), indicating that structural context captures complementary discriminative signals not visible in the sequence representation.

**Fig. 3.**
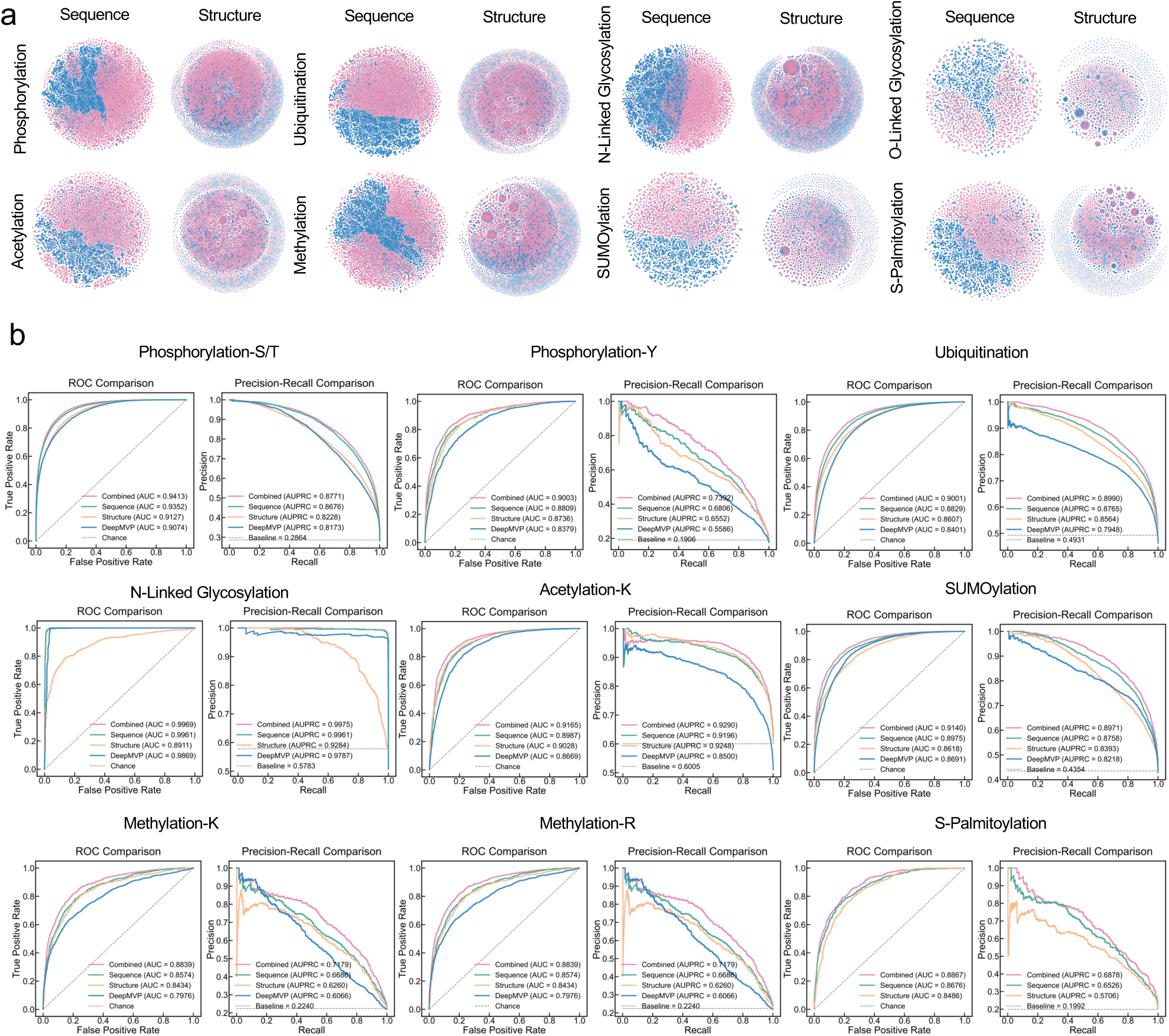
Post-translational modification (PTM) site prediction across multiple modification types. (a) t-distributed stochastic neighbor embedding (t-SNE) visualization of learned embeddings for eight PTM types: phosphorylation, ubiquitination, N-linked glycosylation, O-linked glycosylation, acetylation, methylation, SUMOylation, and S-palmitoylation. For each modification, sequence-derived (left) and structure-derived (right) feature spaces are shown, with PTM-positive (pink) and background (blue) residues. (b) Receiver operating characteristic (ROC) and precision-recall curves for nine PTM classification tasks. The combined model (pink) is compared against sequence-only (green), structure-only (orange), DeepMVP (blue), and chance baseline (dashed). Area under the ROC curve (AUC) and area under the precision-recall curve (AUPRC) values are indicated in the legends.

We evaluated the predictor across 18 PTM types on an independent test set. The dual-modal model achieves AUROC above 0.87 for all PTM types, exceeding 0.90 for 12 of them (Fig. 3b and Extended Data Fig. 5). Benchmarked against DeepMVP^44^, which supports 8 PTM types, the predictor outperforms across all shared types. Complete 5-fold cross-validation results are provided in Extended Data Fig. 6.

To assess whether the learned PTM representations generalize beyond the training proteome, we applied the dual-modal model to proteins associated with 30 major human diseases spanning cancers, neurological disorders, metabolic diseases, and immune conditions (Fig. 4a). Prediction accuracy exceeded 0.80 across the majority of disease-PTM combinations, with glycosylation, phosphorylation, SUMOylation, and ubiquitination reaching near-perfect accuracy in almost all disease categories. Acetylation and methylation showed lower accuracy (0.65–0.80) and greater variability, consistent with their diverse sequence contexts and underrepresentation in current databases. The overall uniformity across 30 diseases indicates that the model captures general sequence-structure features of modification sites rather than family- or tissue-specific patterns.

**Fig. 4.**
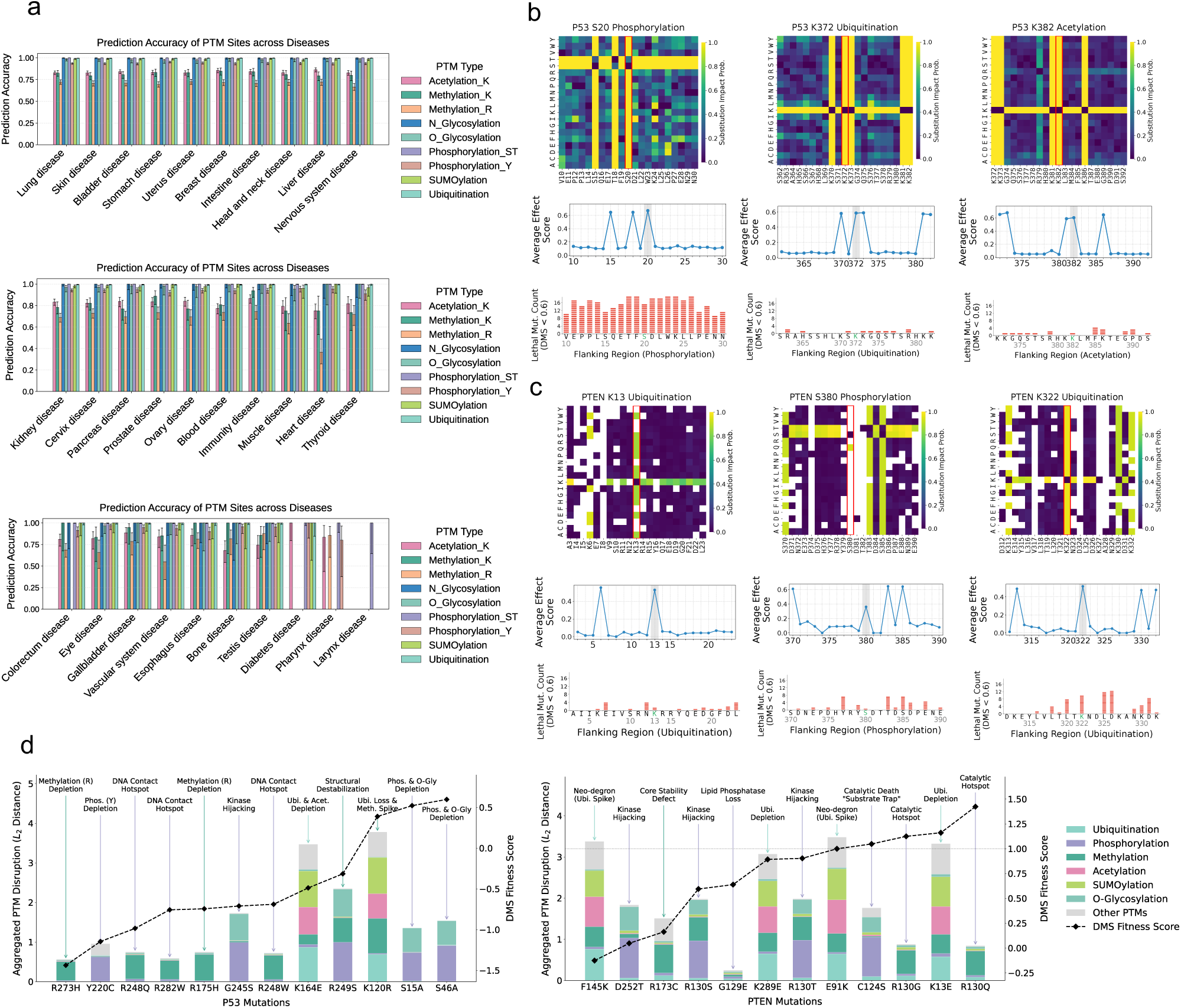
Post-translational modification (PTM) site prediction generalizes across disease-associated proteomes and reveals mutation-driven regulatory disruption. (a) Prediction accuracy of nine PTM types evaluated on proteins associated with 30 major human diseases, grouped across three panels by organ system. Bars represent mean accuracy; error bars indicate standard deviation across proteins within each disease category. (b) In silico saturation mutagenesis of three PTM sites on P53 (S20 phosphorylation, K372 ubiquitination and K382 acetylation). Top, substitution impact probability heatmaps showing predicted PTM disruption for all single-amino-acid substitutions across the flanking region. Middle, position-wise average effect scores. Bottom, number of lethal mutations (deep mutational scanning (DMS) fitness score *<* 0.6) per flanking position. (c) Same analysis as in **b**, applied to three PTM sites on PTEN (K13 ubiquitination, S380 phosphorylation and K322 ubiquitination). (d) Aggregated PTM disruption profiles for disease-associated mutations in P53 (left) and PTEN (right). Stacked bars quantify the *L*_2_ distance between wild-type and mutant PTM probability vectors, decomposed by modification type. Black dashed lines denote experimentally measured DMS fitness scores. Annotations indicate putative molecular mechanisms underlying each mutation’s pathogenic effect.

Beyond site-level classification, we tested whether the PTM predictor captures functional constraints imposed by flanking residues. We focused on six well-studied modification sites on two tumor suppressors, P53 and phosphatase and tensin homolog (PTEN). The P53 sites include S20 phosphorylation, which regulates Mdm2-mediated degradation^45^, and K372 ubiquitination and acetylation. The PTEN sites include K13 ubiquitination, which controls nuclear import ^46^, S380 phosphorylation, which governs the autoinhibitory closed-to-open transition^47,48^, and K322 ubiquitination. At each site, we substituted every position within a 21-residue window with all 20 amino acids and recorded the predicted change in modification probability (Fig. 4b,c). The model assigned the highest disruption scores to the modified residue itself and to flanking positions that form part of the enzyme recognition motif, and these positions were enriched for experimentally lethal mutations identified by deep mutational scanning (DMS) of P53^49,50^ and PTEN^51,52^ (fitness score *<* 0.6). To assess whether mutation effects propagate beyond the local site, we aggregated predicted disruption across all modification types for a panel of cancer-associated missense variants and computed the *L*_2_ distance from the wild-type PTM profile(Fig. 4d). This whole-protein disruption score correlated with DMS fitness for both P53 and PTEN, yet mutations with similar fitness defects showed distinct disruption patterns. For example, DNA-contact mutations in P53 such as R273H and R248W primarily lost methylation, while others gained phosphorylation or shifted ubiquitination, indicating that similar functional outcomes can arise through different changes to the modification landscape^53^.

## 5 Modeling PTM-driven conformational dynamics

We incorporated PTM information as conditioning signals into the dynamics generator to enable modeling of modification-induced conformational changes. This requires MD trajectories paired with explicit PTM annotations for fine-tuning, yet existing datasets such as ATLAS and mdCATH lack systematic modification labels. To address this gap, we constructed DynaMo-phos, a benchmark dataset of phosphorylated protein MD trajectories. We selected phosphorylation as the initial PTM type owing to its prevalence in eukaryotic proteomes. Phosphorylation benefits from extensive experimentally validated site annotations in PhosphoSitePlus ^33^ and mature residue parameterization (SEP, TPO, and PTR) in the CHARMM36m force field, ensuring physically reliable simulations. All simulations were performed as all-atom MD under unified CHARMM36m conditions^54^. The final dataset comprises 580 phosphorylated systems covering phosphoserine (SEP), phosphothreonine (TPO), and phosphotyrosine (PTR) modification types, with a cumulative simulation time of 12.24 *µ*s and diverse structural topologies (Extended Data Fig. 7c). The ProteinFlux architecture is agnostic to modification type and can be extended to additional PTMs as validated force-field parameters become available.

To isolate the contribution of each training stage and the PTM conditioning mechanism, we evaluated three model variants. ProteinFlux-Pretrain is the base model trained only on unmodified protein trajectories, with no exposure to PTM data, serving as a lower bound that reflects general dynamics knowledge. ProteinFlux-Uncond is fine-tuned on DynaMo-phos trajectories but receives no PTM embedding as input, testing whether exposure to phosphorylated dynamics data alone is sufficient to capture PTM-induced conformational changes. ProteinFlux-PTM is the full model, fine-tuned on DynaMo-phos with residue-level PTM embeddings provided as explicit conditioning signals through the adapter module.

Across all three phosphorylation types, ProteinFlux-PTM consistently achieves the highest per-residue RMSF Pearson correlation, followed by ProteinFlux-Uncond and then ProteinFlux-Pretrain (Fig. 5e), confirming that PTM conditioning captures modification-specific flexibility patterns beyond what pre-training alone provides. We further performed an embedding strategy ablation on DynaMo-phos, comparing variants using amino acid embeddings (AA emb), ESM2 evolutionary embeddings, and full PTM conditioning in terms of RMSD coverage and RMSF correlation (Extended Data Fig. 7b). These results revealed a key finding. ESM2 evolutionary embeddings are a necessary condition for generalizing dynamics prediction across protein families, and PTM-specific semantic conditioning on top of ESM2 further substantially improves the prediction accuracy of phosphorylation-induced conformational changes. Neither component alone is sufficient. ESM2 provides the evolutionary foundation for cross-family generalization, while PTM conditioning encodes modification-specific effects upon this foundation.

**Fig. 5.**
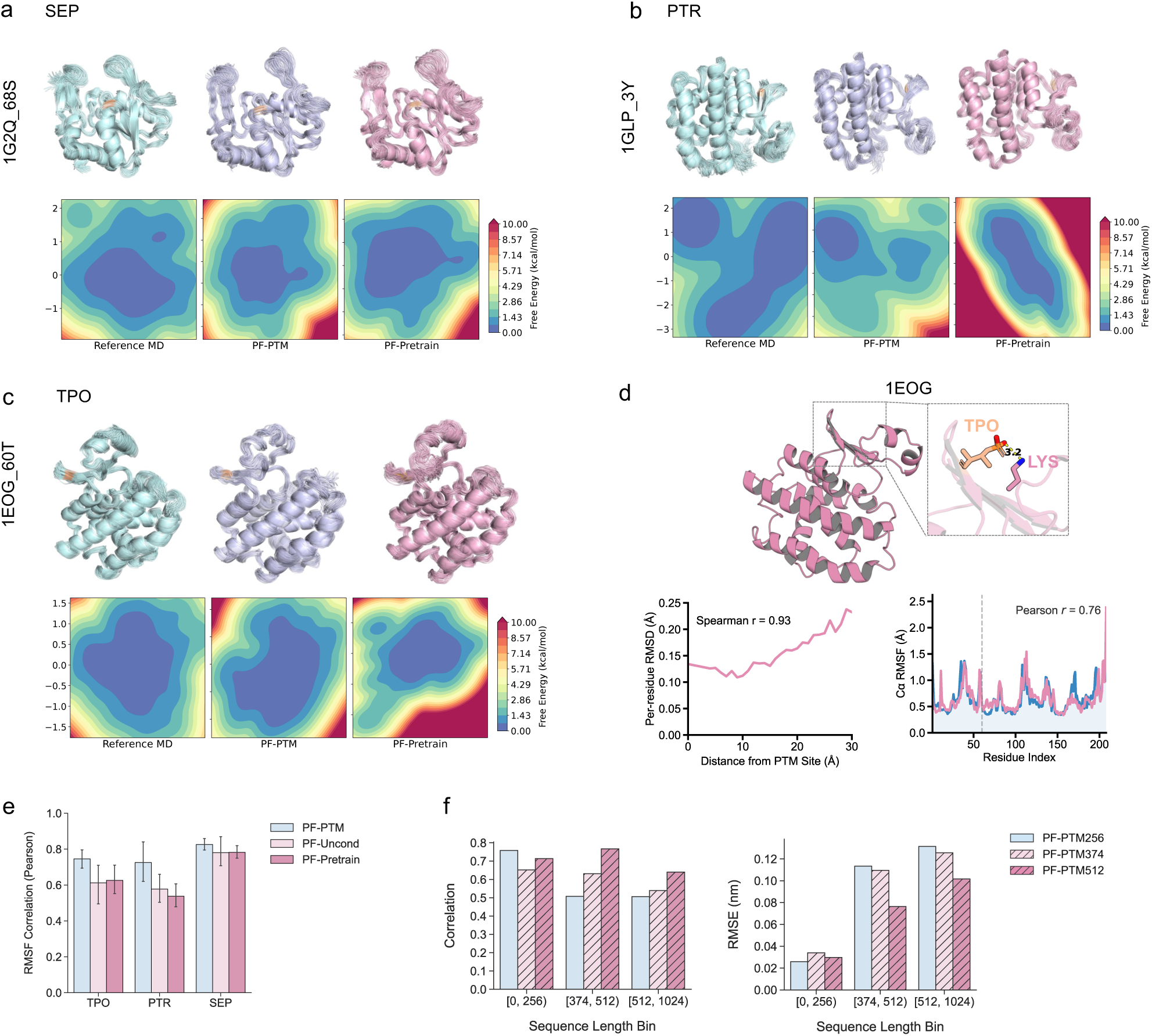
Post-translational modification (PTM)-conditioned dynamics generation. (a-c) Representative case studies for three phosphorylation types: (a) SEP (PDB: 1G2Q), (b) PTR (PDB: 1GLP), and (c) TPO (PDB: 1EOG). Top: superimposed ensembles from reference MD (teal), PF-PTM (purple), and PF-Pretrain (pink). Bottom: free energy landscapes (kcal/mol) projected onto the first two principal components (PC1, PC2). (d) Detailed analysis of 1EOG. Top: structural overlay of reference MD (teal) and PF-PTM (pink) ensembles, with inset showing the TPO60–LYS interaction (3.2 Å). Bottom left: per-residue RMSD as a function of distance from the PTM site. Bottom right: C*_α_* RMSF profiles for reference MD (blue) and PF-PTM (pink); shaded regions indicate standard deviation. (e) Per-residue root mean square fluctuation (RMSF) Pearson correlation between generated and reference molecular dynamics (MD) ensembles for three model variants: ProteinFlux-PTM (PF-PTM), ProteinFlux-Uncond (PF-Uncond), and ProteinFlux-Pretrain (PF-Pretrain), evaluated separately for TPO, PTR, and SEP systems. (f) Effect of crop length on PF-PTM performance. RMSF correlation (left, higher is better) and root mean square error (RMSE, right, lower is better) are shown for three crop lengths (256, 374, and 512 residues) across sequence length bins.

To determine the optimal crop length for PTM conditioning, we fine-tuned ProteinFlux-PTM with three crop lengths (256, 374, and 512 residues) and compared performance across sequence length bins (Fig. 5f). For proteins under 256 residues, all three variants perform comparably, achieving correlations of approximately 0.70 and RMSE below 0.04 nm. Performance diverges for longer sequences. In the 374–512 residue range, PF-PTM374 and PF-PTM512 outperform PF-PTM256, with PF-PTM512 achieving the highest correlation. The advantage of larger crop lengths becomes most pronounced for proteins exceeding 512 residues, where PF-PTM512 maintains a correlation above 0.65 while PF-PTM256 drops substantially. These results demonstrate that larger crop lengths enable the model to capture long-range conformational effects of PTMs in larger proteins.

Beyond aggregate metrics, we examined whether ProteinFlux-PTM reproduces the conformational landscape of individual phosphorylated proteins. For three representative systems, each carrying a different phosphorylation type (phosphoserine in 1G2Q, phosphotyrosine in 1GLP, and phosphothreonine in 1EOG), we compared free energy surfaces projected onto the first two principal components across reference MD, ProteinFlux-PTM, and ProteinFlux-Pretrain (Fig. 5a–c). In all three cases, ProteinFlux-PTM captures the major free energy basins observed in reference MD simulations, including their relative positions and depths. ProteinFlux-Pretrain recovers the overall topology of the landscape but shows noticeable shifts in basin positions and broader distributions. These results indicate that PTM conditioning is important for reproducing not only residue-level flexibility but also the global thermodynamic landscape shaped by phosphorylation.

Detailed analysis of 1EOG further validates the physical plausibility of the model (Fig. 5d). ProteinFlux-PTM positions phosphothreonine 60 within salt-bridge distance (3.2 Å) of a neighboring lysine residue, a physically plausible electrostatic interaction that may stabilize the local conformation upon phosphorylation. More broadly, per-residue RMSD between ProteinFlux-PTM and reference MD increases smoothly with distance from the PTM site (Spearman *r* = 0.93), indicating that the model achieves the highest prediction accuracy near the modification site, where phosphorylation exerts its most direct structural effects. At the residue level, C*_α_*-RMSF profiles closely match reference MD (Pearson *r* = 0.76), confirming that ProteinFlux-PTM reproduces not only the spatial locality of PTM effects but also the magnitude of dynamic fluctuations across the entire protein.

## 6 Applying ProteinFlux to an out-of-distribution test case: the STREX variant of the BK potassium ion channel

The STREX variant of the large conductance calcium- and voltage-activated potassium (BK) channel is a well-characterised model system for studying how palmitoylation- phosphorylation cross-talk regulates ion-channel function. Palmitoylation anchors the alternatively spliced STREX domain to the inner leaflet of the plasma membrane, whereas PKA-mediated serine phosphorylation causes STREX to dissociate from the membrane, consequently suppressing channel activity ^40^. How phosphorylation drives this membrane-dissociation process at the conformational level has not been described at atomic resolution.

We validated FluxSite on an out-of-distribution test case using the full-length BK channel STREX variant (Fig. 6e). Within the 59-residue alternatively spliced STREX insert (residues 633–691), the module identified Ser636 as the sole residue exceeding the prediction threshold. This corresponds to the experimentally validated PKA phosphorylation site, whose phosphorylation mediates STREX-variant-specific channel inhibition^38^. Beyond identifying the correct site, we sought to understand what structural features drive the model’s prediction. Attention weight analysis reveals that the model assigns the highest weight to Ser636 and its spatially neighboring residues within the prediction window (Fig. 6f,g). These high-attention residues cluster in three-dimensional space, as shown by the pairwise C*_α_* distance matrix, suggesting that the model captures structural context relevant to kinase recognition in addition to linear sequence motifs. Outside the insertion, the module identified additional experimentally characterised regulatory residues, including Ser700 (probability 0.97), a known PKC-dependent phosphorylation site ^41^, and Ser927 (probability 0.97), the central PKA-dependent activation site within the canonical RQPS motif^39^. Extended Data Fig. 9 further visualizes the attention patterns and local spatial proximity for the phosphorylation sites Ser700 and Ser927 and the palmitoylation sites Cys645 and Cys646.

**Fig. 6.**
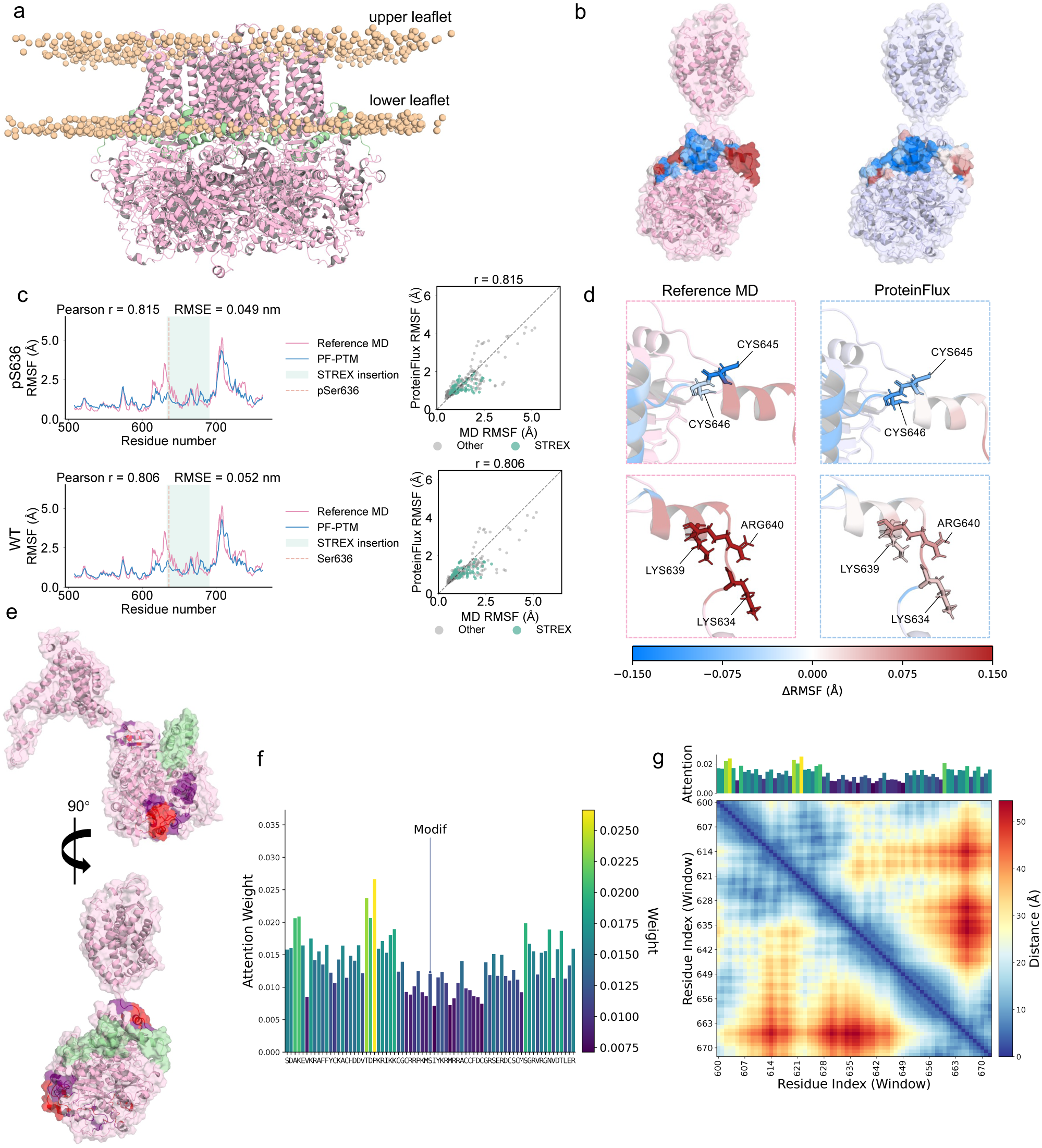
Predicting phosphorylation-induced conformational changes in the BK channel STREX domain. (a) BK channel tetramer embedded in a lipid bilayer, with the STREX insertion highlighted in green. (b) Per-residue ΔRMSF (phosphorylated − unmodified) mapped onto the molecular surface of a single monomer for reference MD (left) and ProteinFlux (right). Blue, reduced flexibility; white, no change; red, increased flexibility. (c) C*_α_* RMSF profiles for the 256-residue BK fragment (residues 508–763) comparing reference MD (pink) and ProteinFlux (blue) under phosphorylated (pSer636, top) and unmodified (WT, bottom) conditions. The STREX region (residues 633–691) is shaded green; dashed line marks Ser636. Right panels show corresponding scatter plots, with STREX residues in green. (d) ΔRMSF mapped onto key functional residues for reference MD (left, pink box) and ProteinFlux (right, blue box), including palmitoylation sites Cys645/Cys646 (upper) and polybasic residues Lys634/Lys639/Arg640 (lower). (e) Predicted phosphorylation sites mapped onto the full BK monomer from two orientations, colored by prediction probability (red, high; purple, moderate). (f) Per-residue attention weights from the site-prediction module for the local sequence window surrounding predicted site Ser636. (g) Joint visualization of attention weights (top) and pairwise C*_α_* distance matrix (bottom) within the STREX prediction window.

To evaluate the dynamics-generation module, we extracted a 256-residue fragment centred on Ser636 (residues 508–763) and generated conformational ensembles under both unmodified and phosphorylated conditions. Dynamics results for pSer636, pSer700 and pSer927 are shown in Extended Data Fig. 8. As ProteinFlux generates trajectories without explicit membrane representation, we focused on local conformational dynamics and adopted RMSF profiles as the primary readout, validated against independent all-atom MD simulations (Fig. 6c). Under the phosphorylated condition, the generated RMSF profile shows strong agreement with the MD reference across the entire fragment (Pearson *r* = 0.815, RMSE = 0.049 nm). Under the unmodified condition, agreement remains robust (Pearson *r* = 0.757, RMSE = 0.047 nm). In both conditions, the model captures the elevated flexibility within the alternatively spliced STREX insertion (residues 633–691), consistent with structural disorder of this domain. RMSF agreement decreases near the fragment boundaries, consistent with the crop-length effect characterised in Fig. 5e, where residues at the edges lack complete structural context.

To assess whether ProteinFlux captures the conformational consequences of phosphorylation, we computed the per-residue change in flexibility (ΔRMSF) and compared the resulting profiles between MD and ProteinFlux (Fig. 6b,d). Within the STREX insertion, basic residues immediately flanking the phosphorylation site, Lys634, Lys639 and Arg640, show increased flexibility upon phosphorylation in both MD and ProteinFlux predictions. This is consistent with the proposal that introduction of negative charge at Ser636 destabilises the polybasic domain that serves as a membrane-association surface ^55^. In contrast, the palmitoylation sites Cys645 and Cys646, located nine residues downstream of the phosphorylation site, show reduced flexibility in both MD and ProteinFlux. Since ProteinFlux predictions were generated without membrane representation, this convergent observation indicates that conformational restriction at the palmitoylation sites arises at least in part from an intrinsic intramolecular response to phosphorylation, independent of changes in membrane proximity. This case study validates ProteinFlux’s integrated pipeline on a biologically relevant system. The site-prediction module accurately identifies validated phosphorylation sites without prior annotations, and the dynamics-generation module produces conformational trajectories that reproduce MD-derived flexibility patterns and capture the differential effects of phosphorylation on functionally distinct residues within the STREX domain.

## 7 Discussion

The ability to predict how post-translational modifications reshape protein conformational dynamics has been limited by two intertwined challenges. The first is encoding modification-site semantics in a form suitable for conditional generation. The second is obtaining paired training data that links PTM annotations to dynamic trajectories. ProteinFlux addresses both challenges through a unified framework that distills evolutionary information from protein language models into task-specific functional representations and couples these representations with a purpose-built phosphorylation dynamics benchmark.

Our results establish three key findings. Evolutionary information, captured through ESM2 embeddings, is a necessary condition for generalizing dynamics prediction across protein families. On this foundation, PTM-specific semantic conditioning, derived by distilling language model features through supervised site prediction, substantially improves the accuracy of predicted phosphorylation-induced conformational changes. The biological relevance of these predictions is supported by zero-shot application to the BK channel STREX domain, where predicted conformational responses at evolutionarily conserved phosphorylation sites are consistent with their established regulatory roles. Importantly, ProteinFlux generates temporally ordered trajectories rather than unordered conformational snapshots, preserving the sequential structure of conformational transitions and enabling analysis of transition pathways and time-dependent flexibility changes.

These findings suggest a principle that extends beyond PTM-conditioned dynamics. Evolutionary constraints encode the functional conformational logic of proteins, and making this information explicit is critical for accurate conditional dynamics modeling. Generic protein language model embeddings alone prove insufficient for effective conditioning. Rather, the supervised distillation of these embeddings into a task-specific functional semantic space yields intermediate representations that serve as the essential bridge between functional annotation and conformational sampling. This strategy may generalize to dynamics prediction conditioned on other biological signals, including disease-causing mutations and protein–protein interactions.

Several limitations define the current boundaries of this work. DynaMo-phos relies on MD simulation trajectories as reference standards, which are constrained by force-field accuracy and sampling sufficiency. For large-scale allosteric transitions operating on millisecond or longer timescales, the current nanosecond-scale simulations may not achieve adequate conformational sampling. The dataset is currently dominated by phosphorylation, and expansion to modifications such as palmitoylation and ubiquitination will require stringent quality control of trajectory physical fidelity. The BK channel case study validates consistency with known phenotypes, but prospective wet-laboratory experiments driven by model predictions remain to be performed. Future work will examine the applicability of the evolutionary conditioning framework to additional PTM types including glycosylation, acetylation, and methylation. We will also explore dynamics prediction under multi-PTM site conditions to capture combinatorial regulatory effects. Another direction is to extend the approach to mutation-driven dynamics prediction. Finally, ProteinFlux could serve as an upstream tool for allosteric site identification, leveraging PTM-induced dynamic perturbations to locate distal regulatory regions. As the first publicly available benchmark systematically collecting PTM-modified MD trajectories with unified simulation protocols and complete functional-site annotations, DynaMo-phos also provides a reusable community resource for developing and evaluating the next generation of conditional protein dynamics models.

## Data availability

The DynaMo-phos benchmark dataset is deposited on Zenodo (https://doi.org/10.5281/zenodo.20050328). The ATLAS, mdCATH and MISATO datasets used for model pretraining are publicly available from their original sources.

## Code availability

The source code for ProteinFlux and FluxSite, including pretrained model weights, data processing pipelines and evaluation scripts, are available at https://github.com/Mochimo-mo/ProteinFlux.

## Acknowledgements

We thank Lingxuan Liu, Xi Wang, Zhuying Chen, Haocheng Tang, Liwei Wang, and Sentao Zhang for their contributions to the molecular dynamics simulations used to construct the DynaMo-phos dataset. Y. Chi, Q.T. Qian, R. Fu, and Y. Qu are members of the SHENG AI Hub (AI for Life-Course Epidemiology of Stress, Infection, and Healthy Aging). Y. Chi, J.H. Peng, Y.M. Xiao, and Y. Qu are members of the CLRA Hub (Cross Lung Research Alliance). Q.T. Qian, J.H. Peng, D.G. Ma, K.N. Liu, Y.H. Cheng, Y.Q. Deng, J.J. Zhao, S. Su, Y.F. Yao, Y. Qu, R. Fu, J.L. Liu, M.K. Zhao, Y.M. Xiao, Y.Z. Wu, Q.F. Xu, J.R. Wang, and Y. Chi are members of the Center for Biomedical System & Informatics. We acknowledge the support of the SHENG AI Hub, the CLRA Hub, and the Center for Biomedical System & Informatics.

## Author Contributions

Q.T. Qian, D.G. Ma, and Y.H. Cheng developed and implemented the ProteinFlux model. J.H. Peng and Y.Q. Deng developed and implemented the FluxSite model. Q.T. Qian, D.G. Ma, K.N. Liu, J.J. Zhao, S. Su, R. Fu, Y.Z. Wu, Yijun Wang, Q.F. Xu, and J.R. Wang constructed the DynaMo-phos dataset and performed molecular dynamics data processing. Yong Wang and K.Y. Wang provided technical guidance on molecular dynamics simulations. Q.T. Qian and J.H. Peng performed biological validation. M.J. Shipston provided domain expertise on the BK channel and guided the biological validation. Q.T. Qian, J.H. Peng, D.G. Ma, and K.N. Liu wrote the manuscript. Y.F. Yao, Y. Qu, J.L. Liu, M.K. Zhao, Y.M. Xiao, and D.C. Hay contributed to result interpretation and discussion. Y. Chi, M.J. Shipston, Yong Wang and Y.H. Ke supervised the project. All authors read and contributed to the final manuscript.

## Extended Data

**Fig. 1.**
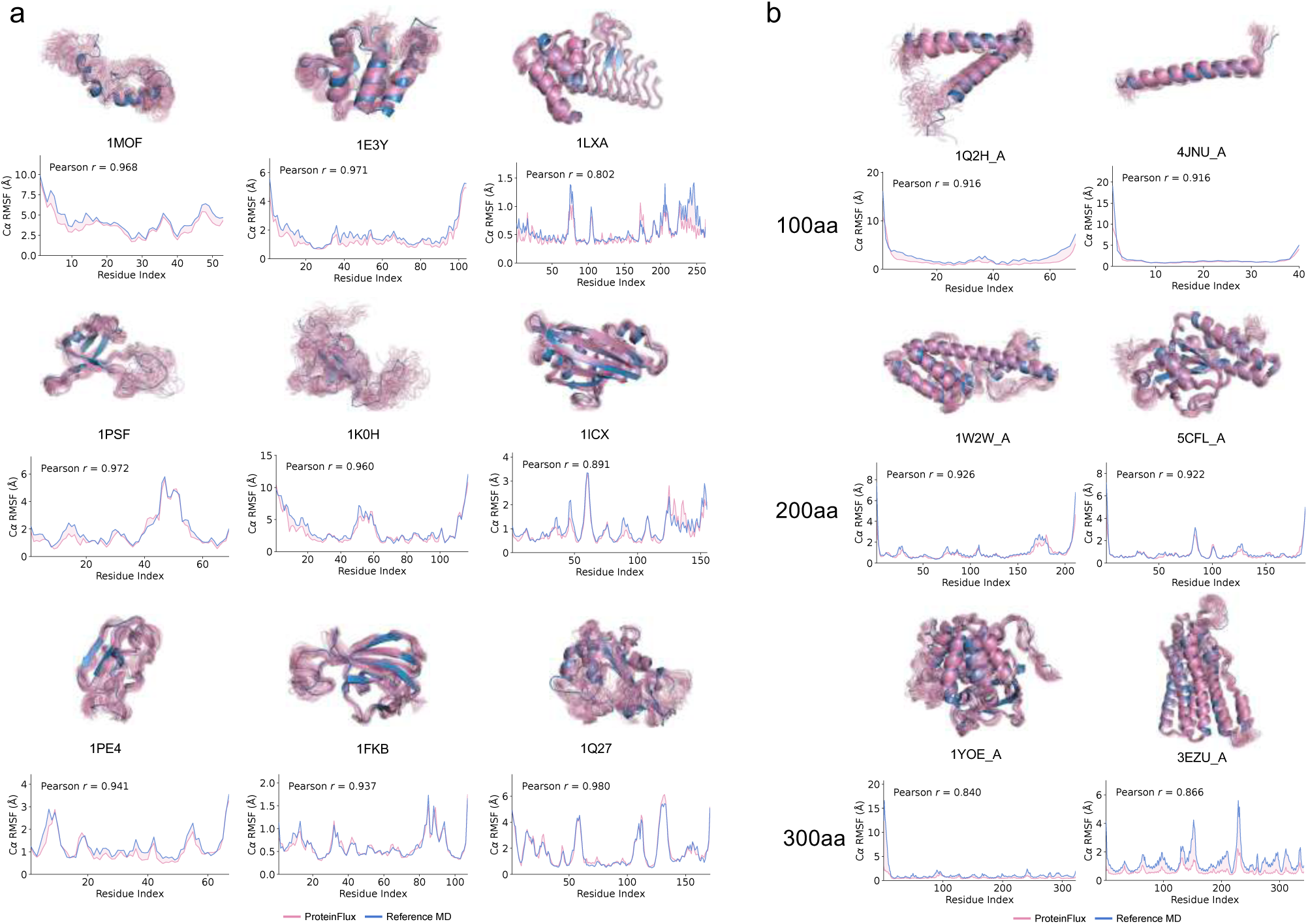
ProteinFlux trained on mixed datasets accurately recapitulates C*_α_* RMSF profiles across diverse protein fold classes and chain lengths. ProteinFlow model trained jointly on MoDEL and ATLAS (100 ps frame interval) was evaluated independently on held-out proteins from both datasets, demonstrating consistent generalization across evaluation sets. (a) Nine proteins from the MoDEL dataset spanning three CATH structural classes: mainly alpha (1MOF, 1E3Y, 1LXA), mainly beta (1PSF, 1K0H, 1ICX), and alpha-beta (1PE4, 1FKB, 1Q27). (b) Six proteins from the ATLAS dataset stratified by chain length: short (*∼*100 aa; 1Q2H A, 4JNU A), medium (*∼*200 aa; 1W2W A, 5CFL A), and long (*∼*300 aa; 1YOE A, 3EZU A).

**Fig. 2.**
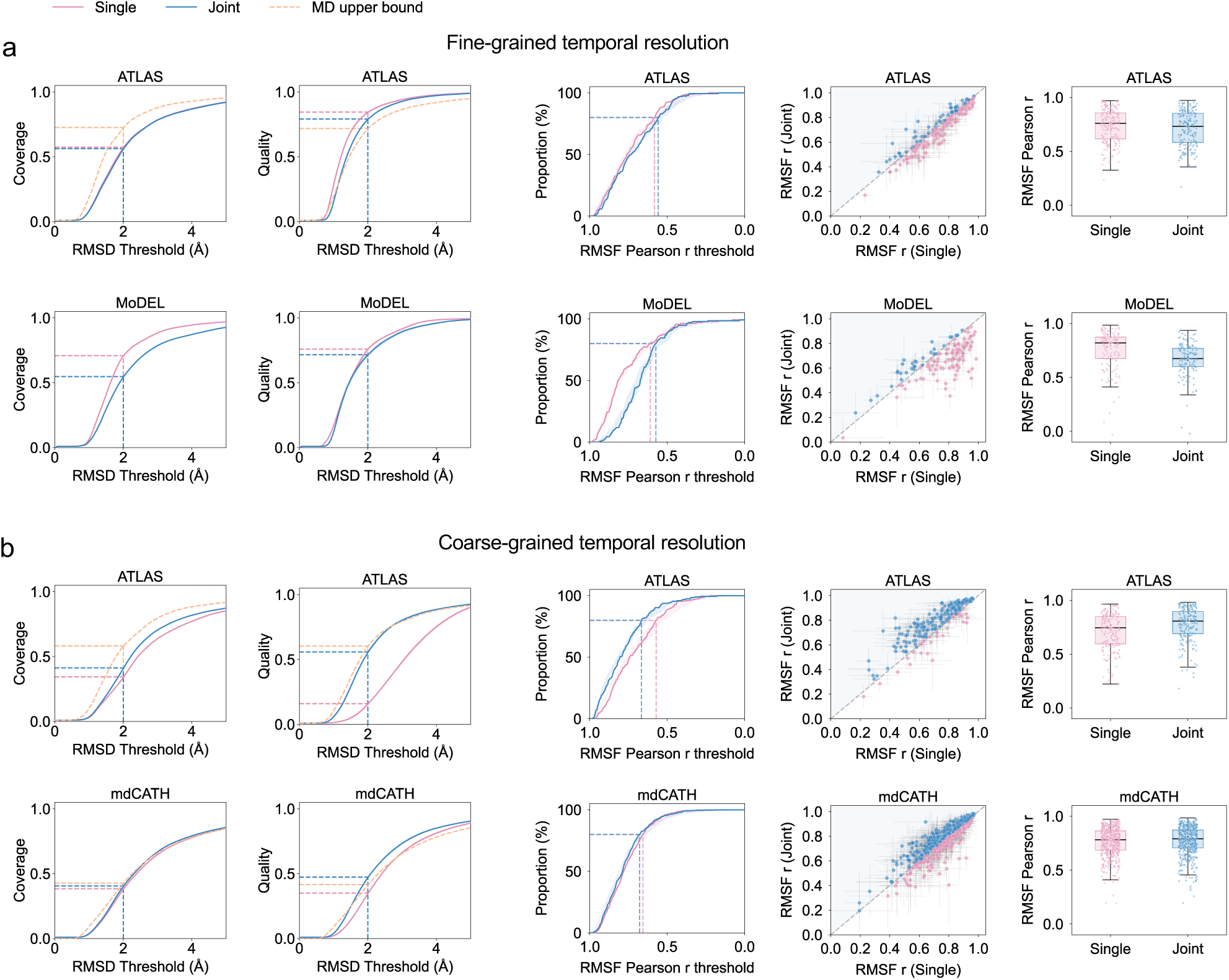
Data strategy ablation: single-dataset (Single) versus multi-dataset joint training (Joint) without ESM2 evolutionary representations. All models use amino acid type embeddings to ensure single-variable control. **a**, Evaluation at fine-grained temporal resolution (100,ps) on ATLAS (top) and MoDEL (bottom) test sets. Joint training degrades performance on ATLAS while yielding marginal improvement on MoDEL, revealing negative transfer between datasets with matched temporal resolution but different simulation protocols. **b**, Evaluation at coarse-grained temporal resolution (1,ns) on ATLAS (top) and mdCATH (bottom) test sets. Joint training similarly impairs ATLAS performance, confirming that sequence-level amino acid embeddings alone are insufficient to reconcile heterogeneous datasets. Columns from left to right: RMSD coverage curves, RMSD quality curves, cumulative distribution of per-protein RMSF Pearson correlation, per-protein RMSF correlation scatter plots, and box plots. Orange dashed lines indicate the MD replica upper bound. Five replica trajectories are generated per protein with different random seeds.

**Fig. 3.**
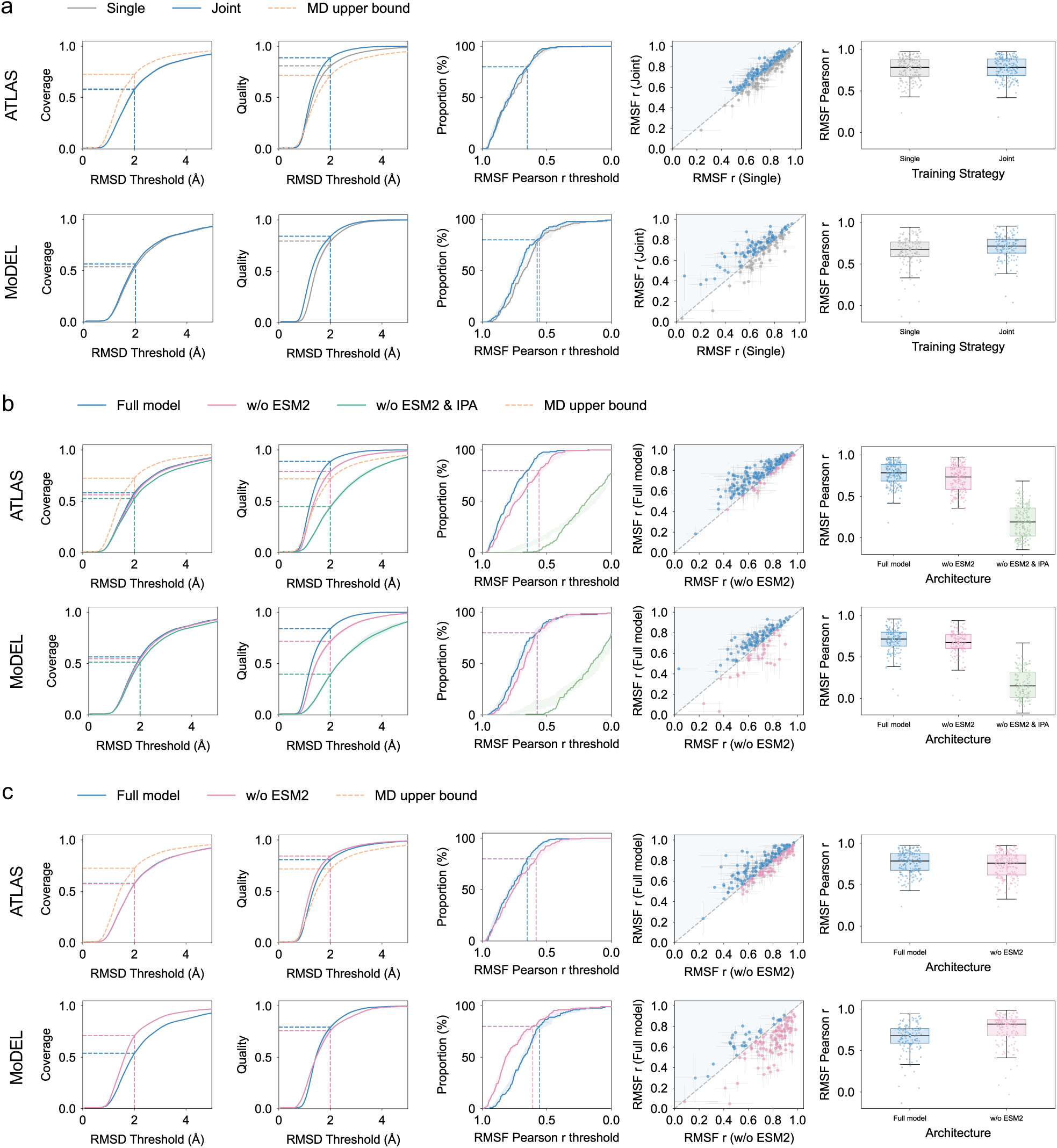
Performance evaluation of training strategies and architectural variants at 100 ps sampling intervals. (a) Comparison of single-dataset (Single) versus multi-dataset joint training (Joint) with ESM2 evolutionary representations, evaluated on ATLAS (top) and MoDEL (bottom) test sets. Joint training substantially improves coverage and RMSF correlation on ATLAS while maintaining comparable performance on MoDEL, indicating that multi-dataset learning does not introduce negative transfer. (b) Architectural ablation under multi-dataset joint training: full model, without ESM2 (w/o ESM2), and without both ESM2 and IPA (w/o ESM2 & IPA). Removing ESM2 leads to marked performance degradation, demonstrating that evolutionary representations are essential for cross-dataset transfer. (c) Comparison of the full model versus without ESM2 under single-dataset training, trained and evaluated on ATLAS (top) and MoDEL (bottom) separately. The contribution of ESM2 is relatively modest in the single-dataset setting, contrasting with its pronounced effect in joint training and revealing an interaction between evolutionary representations and data heterogeneity. Columns from left to right: RMSD coverage curves, RMSD quality curves, cumulative distribution of per-protein RMSF Pearson correlation, per-protein RMSF correlation scatter plots, and box plots. Orange dashed lines indicate the MD replica upper bound.

**Fig. 4.**
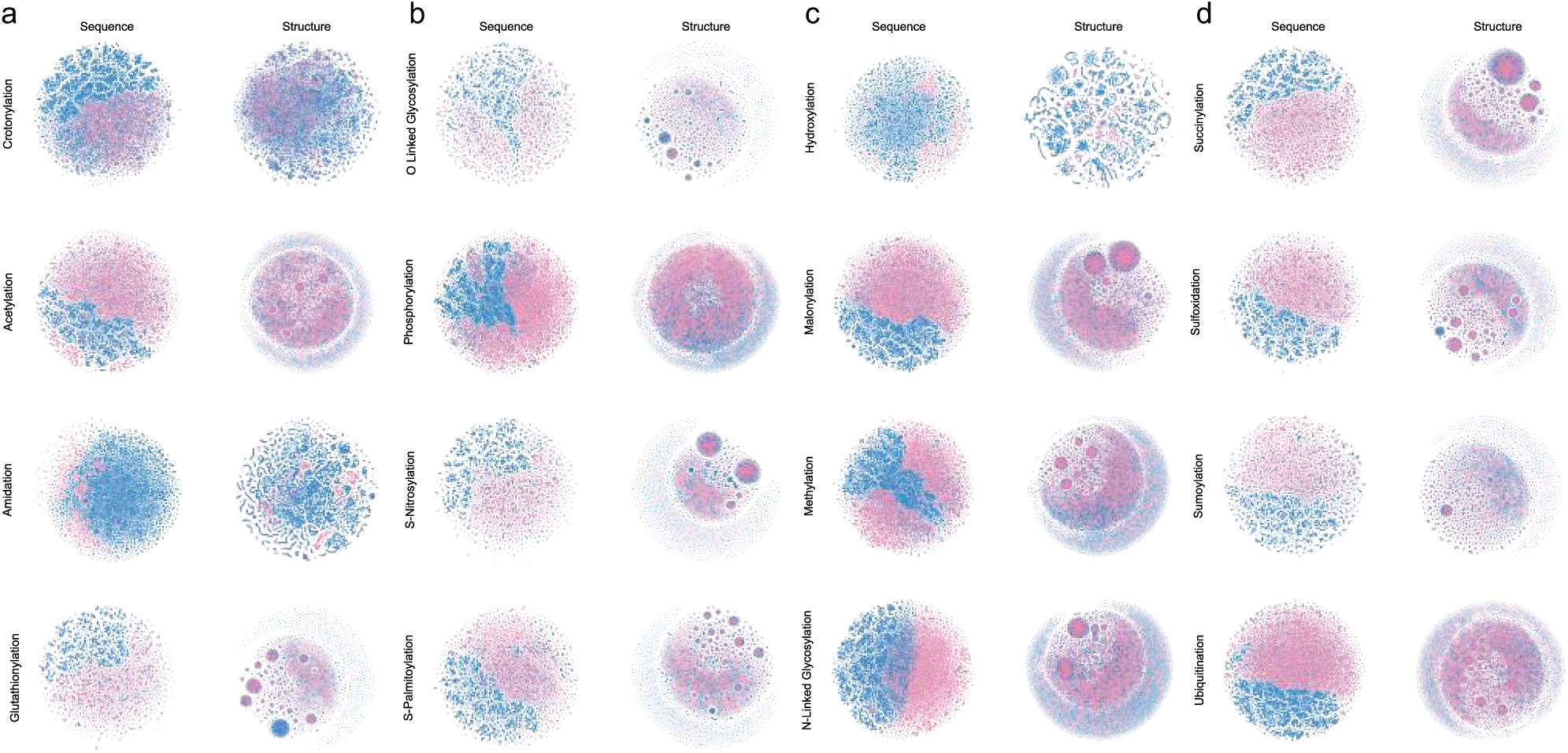
(a-d) t-SNE visualization of sequence and structure features across all 18 PTM types. t-SNE projections of sequence embeddings (left) and structure embeddings (right) for each PTM type. Pink and blue dots represent PTM-positive and PTM-negative sites, respectively. Phosphorylation (Ser/Thr and Tyr) and methylation (Lys and Arg) subtypes are merged.

**Fig. 5.**
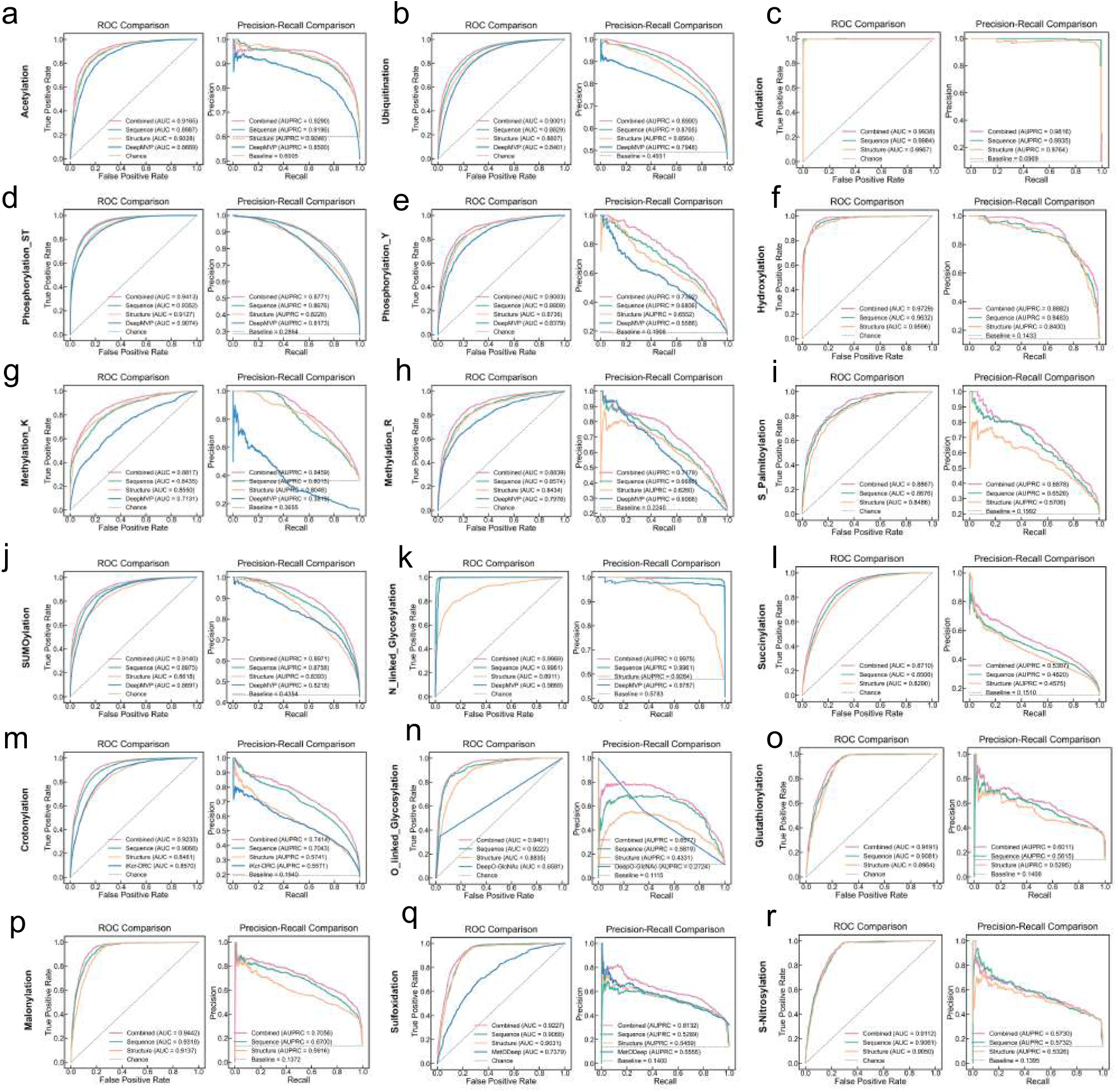
Performance comparison of PTM site prediction across 18 modification types. (a–r) Receiver operating characteristic (ROC) curves (left) and precision-recall (PR) curves (right) evaluating the prediction of 18 distinct PTMs. Predictive performance is compared for the full dual-branch model (Combined; pink), the sequence-only branch (Sequence; blue), the structure-only branch (Structure; teal), and the existing baseline DeepMVP (green). Dashed lines indicate the random chance baseline. The area under the ROC curve (AUROC) and area under the PR curve (AUPRC) values for each approach are provided within the respective panels.

**Fig. 6.**
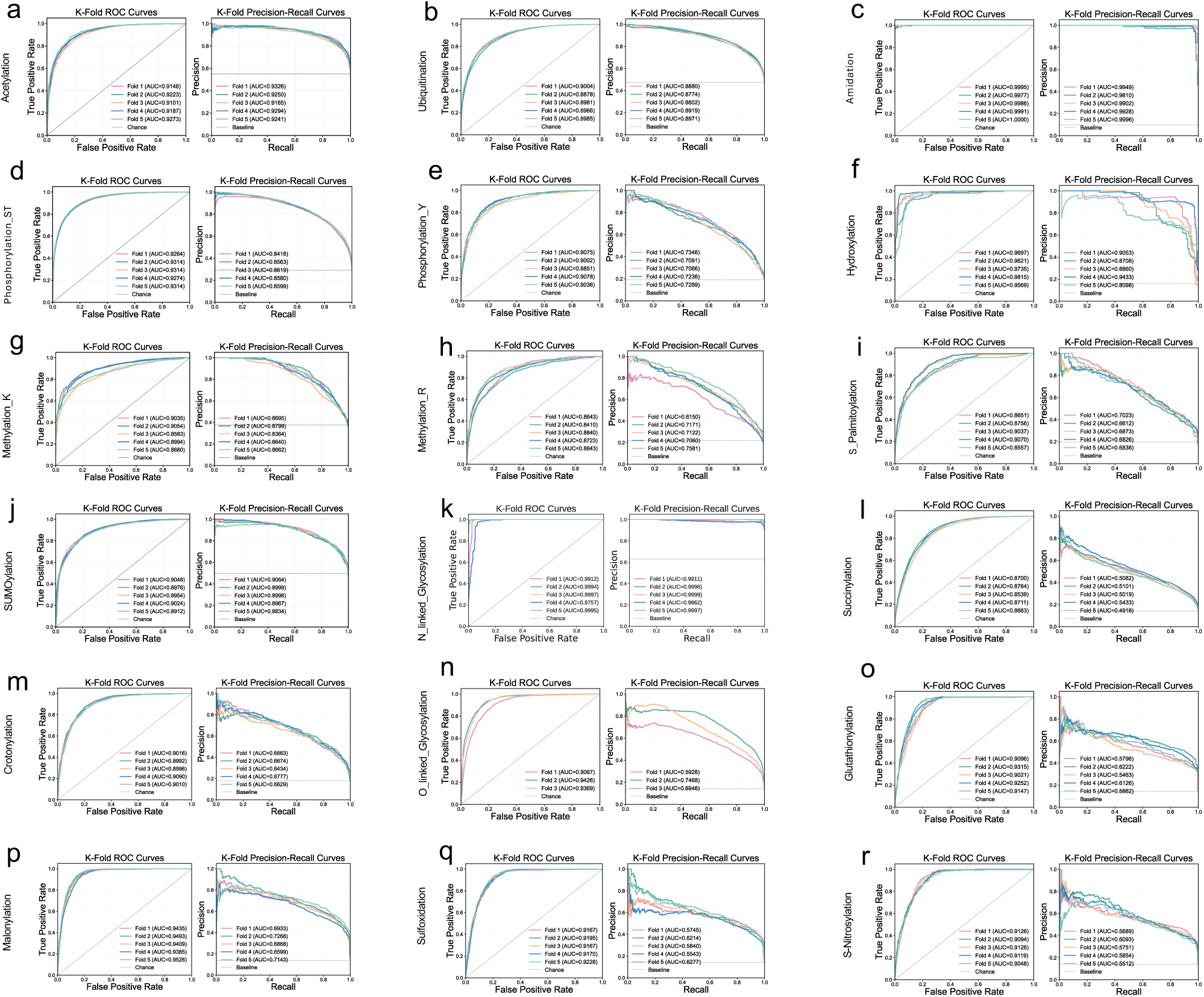
5-fold cross-validation performance of the model across 18 protein post-translational modifications (PTMs). (a–r) 5-fold cross-validation results for 18 distinct types of PTMs. Each sub-figure displays receiver operating characteristic (ROC) curves (left) and precision-recall (PR) curves (right). The five colored curves within each plot illustrate the predictive performance of the model for each individual cross-validation fold (Fold 1 to Fold 5). Values in parentheses within the legends denote the area under the ROC curve (AUC) and the area under the PR curve (AUPRC) for the corresponding fold. The grey diagonal dashed lines in the ROC plots indicate the random chance level, whereas the grey horizontal dashed lines in the PR plots represent the baseline performance.

**Fig. 7.**
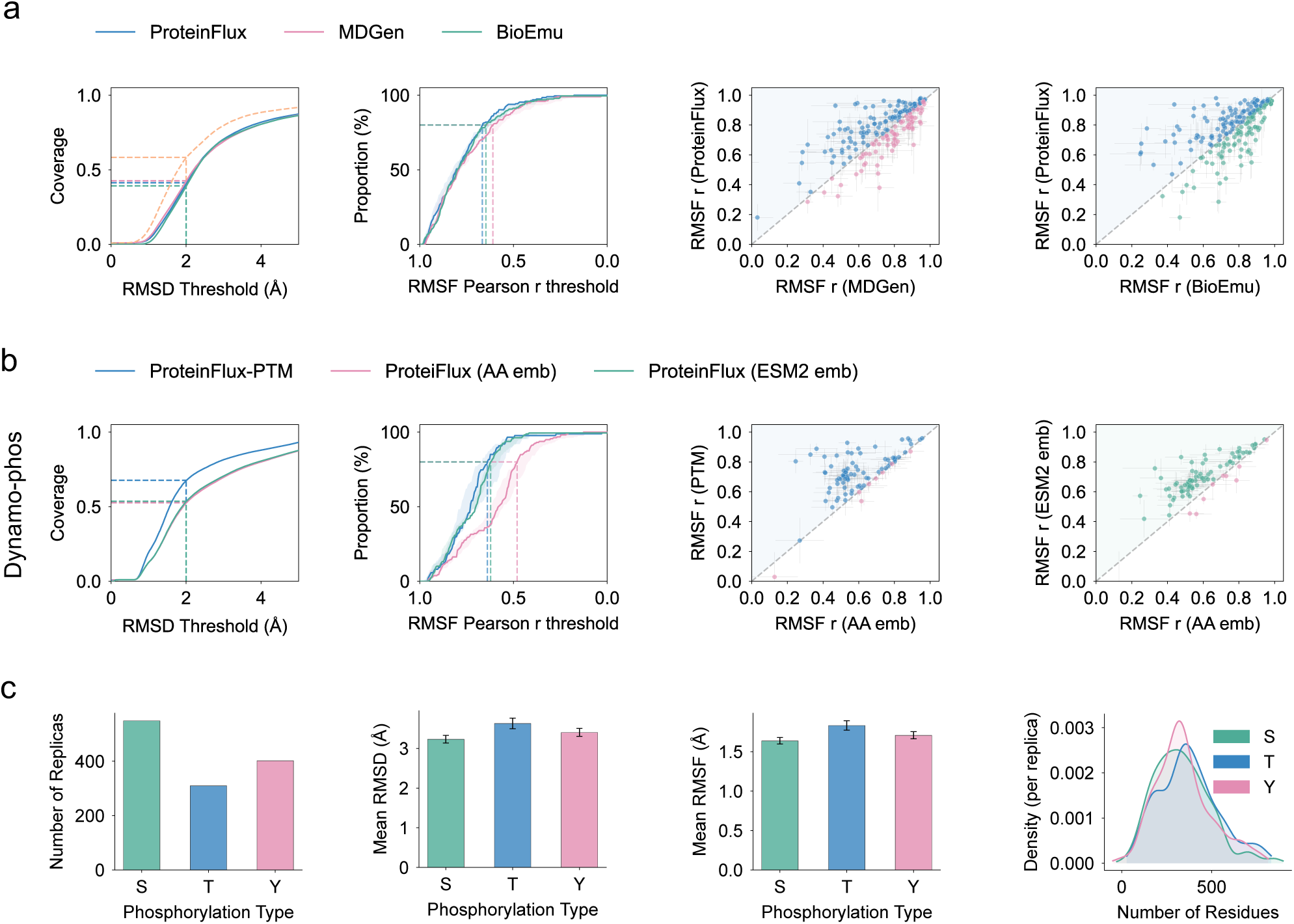
ProteinFlux model evaluation and DynaMo-phos dataset characterization. (a) Comparison of ProteinFlux with baseline methods (MDGen, BioEmu) on the ATLAS test set. From left to right: conformational coverage as a function of RMSD threshold; cumulative proportion of proteins exceeding different RMSF Pearson correlation thresholds; per-protein RMSF correlation scatter plots comparing ProteinFlux against MDGen and BioEmu (diagonal indicates equal performance). (b) Comparison of ProteinFlux-PTM with ablation variants (AA embeddings, ESM2 embeddings) on the DynaMo-phos phosphorylation dataset; panel layout as in **a**. The PTM cross-attention adapter outperforms sequence-embedding-only variants in both RMSD coverage and RMSF correlation, with scatter plots showing systematic improvement of the PTM model over the AA embedding baseline across most proteins. (c) Characterization of the DynaMo-phos dataset by phosphorylation type (serine S, threonine T, tyrosine Y). From left to right: number of trajectories passing quality control; mean RMSD; mean RMSF; density distribution of protein residue counts.

**Fig. 8.**
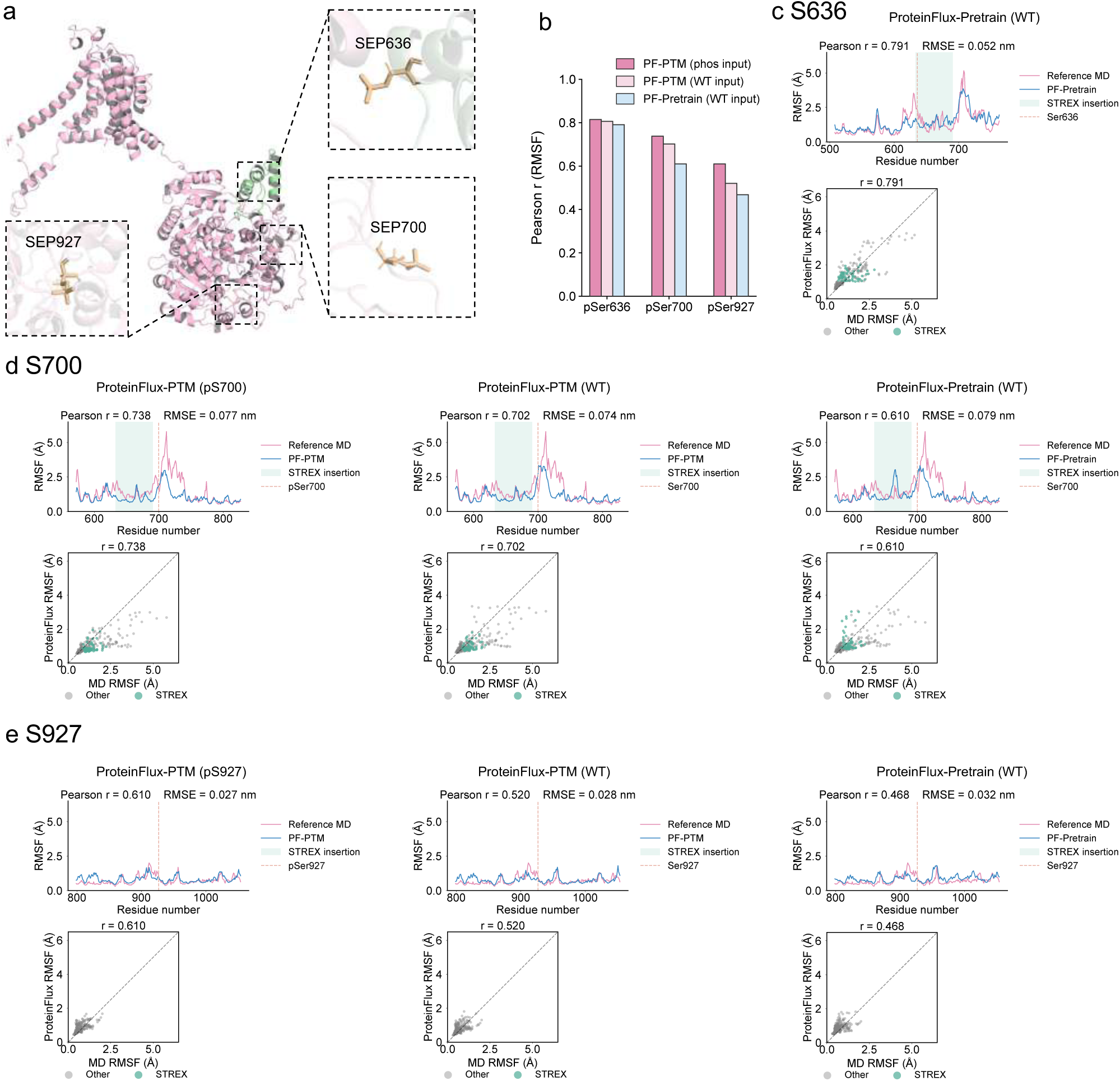
PTM conditioning improves conformational dynamics prediction across multiple BK channel phosphorylation sites. (a) Structural overview of the BK channel monomer, with the STREX insertion highlighted in light green. Three phosphorylation sites pSer636, pSer700, and pSer927 are shown as stick models (orange). Insets show the local structural environment of each site. (b) Summary of RMSF Pearson correlation coefficients across three phosphorylation sites and three inference conditions: ProteinFlux-PTM with phosphorylation input (dark pink), ProteinFlux-PTM with WT input (light pink), and ProteinFlux-Pretrain with WT input (blue). All conditions are evaluated against the corresponding phosphorylated protein MD trajectory as reference. PTM-conditioned inference with phosphorylation input outperforms both WT-input and pretrain baselines across all three sites. (c) Baseline prediction of ProteinFlux-Pretrain for pSer636, shown as per-residue RMSF profiles (top) and scatter plot (bottom); PTM-conditioned results for pSer636 are shown in main figure panel c. (d) Per-residue RMSF profiles (top) and scatter plots (bottom) for pSer700 under three inference conditions, from left to right: ProteinFlux-PTM (phosphorylation input), ProteinFlux-PTM (WT input), and ProteinFlux-Pretrain (WT input). Scatter plot colours distinguish residues within the STREX insertion (green) from all other residues (grey). (e) Per-residue RMSF profiles (top) and scatter plots (bottom) for pSer927 under the same three conditions. pSer927 resides in a disordered C-terminal region distal to the core domain, resulting in lower absolute prediction accuracy compared with the other two sites; nevertheless, PTM-conditioned inference with phosphorylation input yields consistent improvement over both the WT-input and pretrain baselines. All inference results are averaged across ten independent replicas.

**Fig. 9.**
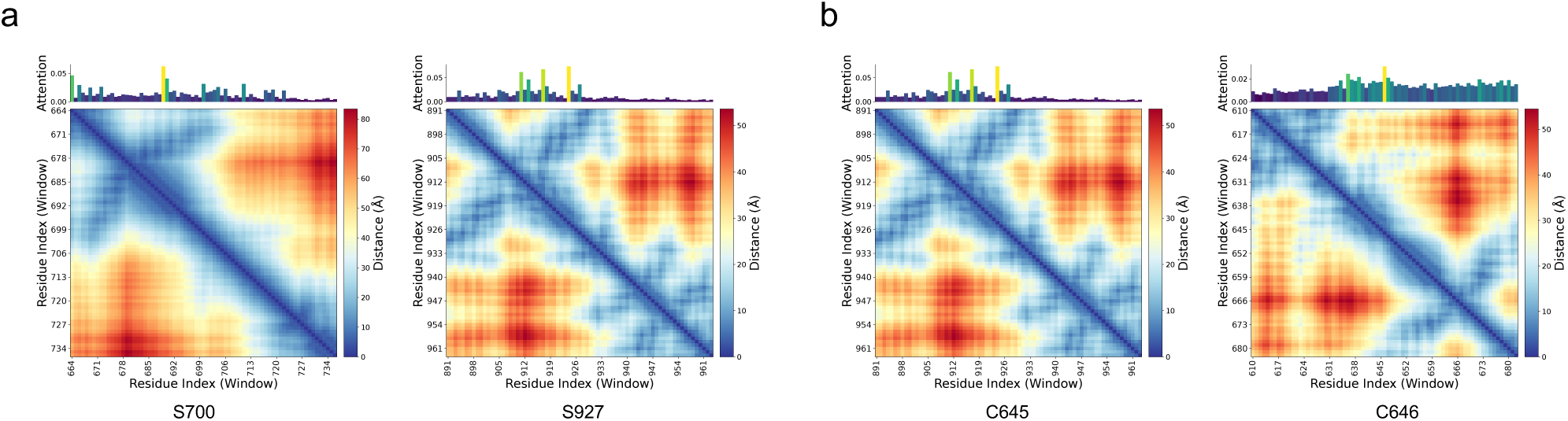
(a) Joint visualization of attention weights and spatial proximity within phosphorylation-site prediction windows (S700 and S927). Prediction windows were centered on Ser700 and Ser927, respectively. For each site, the top bar plot shows per-residue attention weights within the window (colored from low to high), indicating where the model focuses during site identification. The main panel shows the pairwise C*_α_*–C*_α_* distance matrix (Å) for residues within the same window, quantifying their three-dimensional proximity. Residues assigned higher attention tend to cluster spatially, suggesting that the model leverages local structural context in addition to linear sequence information when recognizing phosphorylation-related regulatory sites. (b) Joint visualization of attention weights and spatial proximity within palmitoylation-site prediction windows (C645 and C646). Prediction windows were centered on Cys645 and Cys646, respectively, and attention distributions (top) and C*_α_*–C*_α_* distance matrices (bottom; Å) are shown as in panel a. In the palmitoylation-related region, high-attention residues likewise exhibit pronounced spatial proximity, supporting that the model captures structural features associated with the local environment of lipid-modification sites.

