## Supplementary Information for "ProteinFlux: accurate, rapid and scalable generative prediction of protein dynamics driven by post-translational modifications"

### S.1 Data

#### S.1.1 Pre-training MD Datasets

Pre-training of the base dynamics generator used three publicly available molecular dynamics (MD) simulation repositories: ATLAS[1], MoDEL[2], and mdCATH[3], collectively encompassing 7,781 distinct protein systems and more than 13.3 ms of aggregate simulation time. ATLAS and MoDEL provide all-atom trajectories for complete protein chains; mdCATH supplies domain-level trajectories stratified across the CATH structural hierarchy. Only mdCATH trajectories collected at 320 K were retained for consistency with ATLAS and MoDEL simulation conditions. Summary statistics are in Table S1.

Each residue in every frame is represented by up to 14 heavy-atom coordinates (atom14[4,5]), yielding a four-dimensional array of shape $T \times L \times 14 \times 3$, where $T$ is the frame count and $L$ the sequence length (coordinates in Å). Trajectories are stored in HDF5 format; coordinates are written as `float16` and promoted to `float32` at training time. ATLAS systems contain three independent replicates; mdCATH systems contain five. A single replicate is drawn uniformly at random per system per iteration to ensure diversity without systematic bias.

Non-standard residue types were mapped to canonical counterparts: `CYM/CYX`→`CYS`; `HID/HIE/HIP`→`HIS`; `ASH`→`ASP`; `GLH`→`GLU`; `LYN`→`LYS`; `X`→`GLY`. This convention is applied uniformly in both pre-processing and evaluation pipelines.

During dataset partitioning, each dataset was split independently. For each of ATLAS, MoDEL, and mdCATH, sequence-based clustering was performed using MMseqs2[6] at a 30% sequence identity threshold. Partitioning was carried out at the cluster level: all sequences within a cluster were assigned exclusively to a single split. Multi-replicate systems were treated as a single unit and assigned together. Per-split counts are in Table S2.

**Table S1: Summary statistics of the pre-training MD datasets.**

| **Statistic** | | **MoDEL** | **ATLAS** | **mdCATH** |
|---|---|---|---|---|
| **Systems** | Unique PDB IDs | 1,500 | 1,735 | 4,908 |
| | Total domains | 1,855 | 3,830 | 23,228 |
| **Sequence length (residues)** | Mean | 132.6 | 141.2 | 134.1 |
| | s.d. | 76.3 | 84.5 | 76.4 |
| | Median | 112 | 121 | 112 |
| | Min–Max | 19–539 | 15–759 | 44–487 |
| **CATH** | Superfamilies | 561 | 1,329 | 3,436 |
| **CATH class** | C1 – Mainly $\alpha$ | 469 (25.3%) | 1,039 (27.1%) | 5,834 (25.1%) |
| | C2 – Mainly $\beta$ | 560 (30.2%) | 698 (18.2%) | 5,677 (24.4%) |
| | C3 – Mixed $\alpha/\beta$ | 769 (41.5%) | 1,914 (50.0%) | 10,865 (46.8%) |
| | C4 – Few secondary structures | 55 (3.0%) | 85 (2.2%) | 302 (1.3%) |

**Table S2: Pre-training dataset partitioning.** Cluster-level splits at 30% sequence identity (MMseqs2).

| Dataset | Train | Val | Test | Total |
|---|---|---|---|---|
| MoDEL | 1,200 | 150 | 150 | 1,500 |
| ATLAS | 1,388 | 173 | 174 | 1,735 |
| mdCATH | 3,926 | 490 | 492 | 4,908 |
| **Total** | **6,514** | **813** | **816** | **8,143** |

##### S.1.2 PTM Site Prediction Datasets

**Data Sources and Collection** PTM site annotations were curated from three complementary databases: dbPTM[7], PLMD[8], and UniProt[9], covering 18 modification types: phosphorylation (S/T/Y), ubiquitination, acetylation, SUMOylation, N-glycosylation, O-glycosylation, methylation (K/R), crotonylation, glutathionylation, hydroxylation, malonylation, S-palmitoylation, S-nitrosylation, succinylation, sulfoxidation, and amidation. Structural data were obtained from the AlphaFold Database[4].

**Inclusion and Exclusion Criteria** At the sequence level, sequences with $50 \leq L \leq 1024$ residues were included; longer sequences were truncated around the target PTM site, and sequences containing non-standard amino acid codes (B, J, X, Z) were excluded. At the structure level, residues with pLDDT $< 75$ were masked, and structures with $> 5\%$ missing C$\alpha$ atoms or any C$\alpha$–C$\alpha$ distance $> 4.0$ Å were discarded.

**Safe Negative Sampling** A safe negative sampling protocol was used to address class imbalance. Negatives were drawn strictly from residues with no experimental PTM evidence in dbPTM, excluding "potential" and "ambiguous" annotations, to avoid mislabelling true positives. Hard negatives—sites with high sequence similarity to positives but confirmed as unmodified—were retained to sharpen the decision boundary. Finally, CD-HIT[10] at 30% identity was applied to the negative set to remove redundancy.

**Dataset Splitting** A strict protein-level split was applied (train 80%, validation 10%, test 10%) based on unique UniProt IDs, ensuring that no PTM sites from the same protein appear across splits. A fixed random seed (42) was used for reproducibility. Table S3 details the number of positive and negative samples for each PTM type in both the training and test partitions.

##### S.1.3 DynaMo-phos: Phosphorylated Protein MD Trajectory Dataset

DynaMo-phos is an in-house all-atom MD dataset of phosphorylated proteins constructed to support PTM-conditioned dynamics generation. Source proteins were drawn from the PTM site prediction

**Table S3: Training dataset statistics for PTM site prediction.** Number of positive and negative samples across 18 PTM types in the training and testing sets.

| PTM Type | Target Residue(s) | Training Set | | Test Set | |
|---|---|---|---|---|---|
| | | Positive | Negative | Positive | Negative |
| Phosphorylation | S, T | 164,942 | 448,290 | 17,189 | 44,050 |
| Phosphorylation | Y | 10,397 | 48,369 | 1,023 | 4,792 |
| Ubiquitination | K | 96,133 | 109,355 | 10,228 | 11,866 |
| Acetylation | K | 30,057 | 33,254 | 2,837 | 2,724 |
| Methylation | K | 9,679 | 38,155 | 1,236 | 3,708 |
| Methylation | R | 8,617 | 41,105 | 979 | 3,475 |
| N-linked glycosylation | N | 7,040 | 7,535 | 862 | 838 |
| Sumoylation | K | 35,249 | 36,220 | 3,691 | 4,927 |
| Succinylation | K | 26,907 | 180,705 | 2,961 | 20,799 |
| O-linked glycosylation | S, T | 28,114 | 88,801 | 2,115 | 5,548 |
| S-palmitoylation | C | 5,803 | 14,725 | 592 | 1,506 |
| Malonylation | K | 10,713 | 21,426 | 1,281 | 2,562 |
| S-Nitrosylation | C | 3,744 | 7,488 | 399 | 798 |
| Sulfoxidation | M | 6,752 | 13,504 | 766 | 1,532 |
| Glutathionylation | C | 3,656 | 7,312 | 389 | 778 |
| Amidation | Any | 2,997 | 17,982 | 315 | 1,890 |
| Hydroxylation | Any | 1,826 | 10,956 | 570 | 3,420 |
| Crotonylation | K | 12,263 | 60,101 | 3,344 | 15,010 |

datasets (Section S.1.2). For each protein, three independent MD replicas were performed to enhance conformational sampling and assess trajectory reproducibility.

Each simulated system contains a single phosphorylation site. CHARMM36m force field[11] residue types are used: SEP (phosphoserine), TPO (phosphothreonine), and PTR (phosphotyrosine). Charge assignments and protonation states are defined by the force-field residue topology. Protonation states at pH 7.0 were assigned by PDB2PQR and PROPKA 3.5[12,13], which predicts p$K_a$ values from local electrostatic and hydrogen-bonding environments and assigns histidine tautomers (HSD/HSE/HSP). GROMACS[14] `pdb2gmx` built the topology (TIP3P water[15]) and rebuilt hydrogens per force-field parameters. A cubic periodic-boundary box was constructed with $Na^+$ counterions to neutralise the total charge.

###### S.1.3.1 MD Simulation Protocol

All simulations were performed with GROMACS v2025.2 in four stages, all using Verlet neighbour lists, PME electrostatics, and $r_c = 1.2\,\mathrm{nm}$ cutoffs.

**Energy minimisation.** Steepest descent until maximum force $< 1000\,\mathrm{kJ\,mol^{-1}\,nm^{-1}}$.

**NVT equilibration (100 ps).** 2 fs steps; V-rescale thermostat at 310 K ($\tau = 0.1\,\mathrm{ps}$); heavy-atom position restraints.

**NPT equilibration (200 ps).** Parrinello–Rahman barostat added (isotropic, 1 bar, $\tau = 2$ ps, compressibility $4.5 \times 10^{-5}$ bar$^{-1}$); restraints maintained. Box dimensions and density converge in this stage.

**Production (10 ns).** Restraints released; 2 fs steps; V-rescale (310 K, $\tau = 0.5$ ps); Parrinello–Rahman (1 bar, $\tau = 2$ ps); LINCS hydrogen-bond constraints; dispersion corrections; centre-of-mass motion removed every 100 steps. Coordinates saved every 100 ps (100 frames, excluding $t = 0$); three replicates per system with independent velocity seeds.

###### S.1.3.2 Trajectory Post-processing and Quality Control

After simulation, PBC molecules were unwrapped, the protein was centred in the box, and backbone alignment was applied to remove global rotation and translation. Protein-only trajectories were then exported using MDAnalysis[16]. Quality control was performed at two levels. Hard gates rejected any trajectory that failed to load or exhibited missing phosphorus (P) atoms relative to annotated PTM sites, ensuring that conditioning labels remained consistent with the underlying molecular structure. Trajectories were assessed by C$\alpha$ RMSD, radius of gyration ($R_g$), and per-residue RMSF, and classified into three quality levels: PASS, WARNING, and FAIL. A trajectory was rejected (FAIL) if the equilibrium-phase C$\alpha$ RMSD exceeded a length-dependent threshold (15 Å for $L \leq 200$; 18 Å for $200 < L \leq 400$; 20 Å for $L > 400$), if the temporal drift exceeded 1.5 Å ns$^{-1}$, or if the $R_g$ coefficient of variation exceeded 10%. A WARNING was assigned when the equilibrium-phase C$\alpha$ RMSD exceeded a lower length-dependent threshold (8 Å for $L \leq 200$; 10 Å for $200 < L \leq 400$; 12 Å for $L > 400$), when the equilibrium RMSD relative fluctuation ($\sigma/\mu$) exceeded 0.3, drift exceeded 1.0 Å ns$^{-1}$, or $R_g$ CV exceeded 6%. Only trajectories rated PASS were retained for inclusion in the DynaMo-phos dataset; WARNING and FAIL systems were excluded.

#### S.2 Model Architecture

##### S.2.1 Dynamics Generation Module

###### S.2.1.1 Protein Representation

Given a protein sequence $\mathbf{s}$ and an initial set of rigid-body frames $\mathbf{T}^{(0)} = \{T_i^{(0)}\}_{i=1}^{L}$ derived from the reference structure, the model learns the conditional distribution $p(\mathbf{x} \mid \mathbf{s}, \mathbf{T}^{(0)})$, where $\mathbf{x}$ is a relative-offset trajectory representation. Specifically, the conformational state of residue $i$ at trajectory frame $\tau$ is encoded as:

$$\mathbf{x}_i^\tau = \left(\Delta\mathbf{q}_i^\tau,\ \Delta\mathbf{t}_i^\tau,\ \boldsymbol{\alpha}_i^\tau\right) \in \mathbb{R}^{21}, \tag{S1}$$

where $\Delta\mathbf{q}_i^\tau \in \mathbb{R}^4$ is the unit quaternion rotation offset relative to $T_i^{(0)}$ (with non-negative real part to resolve sign ambiguity), $\Delta\mathbf{t}_i^\tau \in \mathbb{R}^3$ is the corresponding translational offset, and $\boldsymbol{\alpha}_i^\tau \in \mathbb{R}^{14}$ encodes seven torsion angles $(\phi, \psi, \omega, \chi_1\text{–}\chi_4)$ as sine–cosine pairs. By parameterising all quantities as offsets relative to the reference frame rather than in absolute coordinates, the model benefits from a simplified learning target and maintains structural consistency across the generated ensemble.

**Sequence features.** Residue-level embeddings extracted from ESM-2[17] are projected to a hidden dimension $d = 384$ via a learned linear layer.

**Structural features.** Heavy-atom coordinates are extracted using the atom14 scheme[4]; missing atoms are zero-padded with binary validity masks. Per-residue rigid-body frames $T_i^{(0)} = (R_i^{(0)}, \mathbf{t}_i^{(0)}) \in \mathrm{SE}(3)$ are constructed by Gram–Schmidt orthogonalisation of the N–C$\alpha$–C atom triplet. The frame set $\{T_i^{(0)}\}$ serves both as structural conditioning and as the coordinate system for offset computation.

###### S.2.1.2 SE(3)-Equivariant Geometric Backbone

The model stacks $N_{\text{IPA}}$ Invariant Point Attention (IPA) layers[4] as the core geometric backbone. Unlike standard attention mechanisms, IPA explicitly operates on a dual-representation framework, simultaneously updating 1D single-residue representations and 3D backbone frames (rotations and translations). It calculates attention weights by combining standard dot-product sequence attention with the squared Euclidean distances between projected query and key points in the 3D space. By operating strictly within local frame coordinates, IPA enables rigorous $SE(3)$-equivariant geometric message passing. This ensures that the predicted spatial updates are inherently invariant to global rotations and translations of the protein, eliminating the need for structural data augmentation.

###### S.2.1.3 Flow Matching in Relaxed Euclidean Space

Although the protein backbone and sidechains are fundamentally described by $SE(3)$ rigid bodies and torsion angles, operating directly on complex manifolds can be computationally expensive and prone to numerical instability. Instead, we adopt a *relaxed* Euclidean formulation. We flatten the spatial representations for each residue into a continuous Euclidean vector $\mathbf{x} \in \mathbb{R}^{21}$.

We construct a linear probability path between a standard Gaussian prior $\mathbf{x}_0 \sim \mathcal{N}(\mathbf{0}, \mathbf{I})$ and the target data distribution $\mathbf{x}_1$. The intermediate state at time $t \in [0, 1]$ is defined as:

$$\mathbf{x}_t = (1 - t)\mathbf{x}_0 + t\mathbf{x}_1. \tag{S2}$$

This linear path yields a constant target vector field (velocity) $u_t = \mathbf{x}_1 - \mathbf{x}_0$. The training objective matches the neural network prediction $\hat{v}_\theta$ to this ground-truth continuous vector field:

$$\mathcal{L}_{\text{coord}} = \mathbb{E}_{t\sim\mathcal{U}[0,1],\, \mathbf{x}_0\sim\mathcal{N}(\mathbf{0},\mathbf{I}),\, \mathbf{x}_1} \left[ \left\| \hat{v}_\theta(\mathbf{x}_t, t, \mathbf{s}, \mathbf{T}^{(0)}) - (\mathbf{x}_1 - \mathbf{x}_0) \right\|^2 \right]. \tag{S3}$$

To ensure geometric validity during inference, the predicted flattened coordinates can be projected or decoded back into physically valid 3D structures. The mean squared error loss is masked and averaged over the $L$ valid residues across the trajectory length $T$. Algorithm S1 details the full training-step procedure.

**Algorithm S1** Linear Flow Matching Loss Computation

**Require:** Velocity network $v_\theta$; target trajectory $\mathbf{x}_1 \in \mathbb{R}^{T\times L\times 21}$; mask $\mathbf{m} \in \{0, 1\}^{T\times L}$; conditioning $\mathbf{s}$, $\mathbf{T}^{(0)}$.

Sample prior $\mathbf{x}_0 \sim \mathcal{N}(\mathbf{0}, \mathbf{I})$ with the same shape as $\mathbf{x}_1$.
Sample time step $t \sim \mathcal{U}[0, 1]$.
▷ Linear interpolation
$\mathbf{x}_t := (1 - t)\mathbf{x}_0 + t\mathbf{x}_1$.
▷ Target velocity
$u_t := \mathbf{x}_1 - \mathbf{x}_0$.
▷ Network prediction
$\hat{v}_\theta := v_\theta(\mathbf{x}_t, t, \mathbf{s}, \mathbf{T}^{(0)})$.
▷ Masked MSE loss
$\mathcal{L} := \frac{1}{\sum_{i,\tau} m_{i,\tau}} \sum_{i,\tau} m_{i,\tau} \|\hat{v}_\theta^{(i,\tau)} - u_t^{(i,\tau)}\|^2$.
**return** $\mathcal{L}$.

###### S.2.1.4 Inference Pipeline

Generation starts from $\mathbf{x}_0 \sim \mathcal{N}(\mathbf{0}, \mathbf{I})$ and integrates the probability-flow ODE:

$$\frac{\mathrm{d}\mathbf{x}_t}{\mathrm{d}t} = v_\theta(\mathbf{x}_t,\, t,\, \mathbf{s},\, \mathbf{T}^{(0)}), \quad t \in [0, 1]. \tag{S4}$$

Integration uses the Dormand–Prince adaptive solver (`dopri5`)[18], $\varepsilon_{\text{abs}} = 10^{-6}$, $\varepsilon_{\text{rel}} = 10^{-3}$. The generated $\mathbf{x}_1 = (\Delta\mathbf{q}_i, \Delta\mathbf{t}_i, \boldsymbol{\alpha}_i)$ is composed with $\mathbf{T}^{(0)}$ to recover global C$\alpha$ coordinates, followed by torsion-angle decoding to all-atom coordinates (Algorithm S2).

**Algorithm S2** Conformational Ensemble Generation (Inference)

**Require:** Trained $v_\theta$; sequence $\mathbf{s}$; frames $\mathbf{T}^{(0)}$; frame count $T$; tolerances.

$\mathbf{x}_0 \sim \mathcal{N}(\mathbf{0}, \mathbf{I}),\ \ \mathbf{x}_0 \in \mathbb{R}^{T\times L\times 21}$.

Integrate Eq. S4 via `dopri5` to obtain $\mathbf{x}_1$.

**for** $\tau = 1, \ldots, T;\ \ i = 1, \ldots, L$ **do**

$\quad R_i^{(\tau)} {:=} R_i^{(0)} \cdot \mathrm{QuatToRot}(\Delta\mathbf{q}_i^{(\tau)});\quad \mathbf{t}_i^{(\tau)} {:=} \mathbf{t}_i^{(0)} + R_i^{(0)} \Delta\mathbf{t}_i^{(\tau)}$.

**end for**

$\hat{\mathbf{X}}_{\text{all-atom}} {:=} \mathrm{FramesToAtom14}(\{\hat{T}_i^{(\tau)}, \boldsymbol{\alpha}_i^{(\tau)}\})$.

**return** $\hat{\mathbf{X}}_{\text{all-atom}}$.

#### S.2.2 FluxSite: PTM Site Prediction

##### S.2.2.1 Dual-Branch Architecture

FluxSite integrates sequence and structural modalities through a dual-tower architecture that processes each modality independently before fusing their representations.

The **sequence tower** takes as input an $L \times 1315$ feature matrix formed by concatenating ESM-2 embeddings (1280-dim) with one-hot amino-acid encodings (35-dim). These features are first linearly projected, then processed by a multi-scale convolutional block with kernel sizes of 3, 5, and 7 to capture local sequence motifs at varying granularities. The resulting representations are refined by a Transformer encoder and subsequently compressed via multi-head attention pooling with rank $r = 5$.

The **structure tower** operates on $L \times 512$ structural embeddings obtained from ESM-IF1[19]. After linear projection, a distance-aware spatial graph attention network (GAT, $N_h = 4$) encodes local geometric context. This is followed by a Transformer encoder augmented with SpatialGNN residual connections, and the same attention-pooling scheme ($r = 5$) is applied.

Both towers yield a site-centric representation $\mathbf{H} \in \mathbb{R}^{B\times d_h}$ and a contextual feature map $\mathbf{C} \in \mathbb{R}^{B\times L\times d_h}$ with $d_h = 256$, which are subsequently combined by the Gated Fusion Module (Table S4).

**Table S4: Dual-branch architecture comparison.**

| Component | Sequence branch | Structure branch |
|---|---|---|
| Input features | ESM-2 + one-hot | ESM-IF1 |
| Input dim. | $1280 + 35 = 1315$ | 512 |
| Projection | Linear→LN→GELU→Drop | Linear→LN→GELU→Drop |
| Local encoder | Multi-scale CNN (k=3,5,7) | Spatial GAT ($N_h = 4$) |
| Global encoder | Transformer | Transformer + SpatialGNN residual |
| Pooling | Attention pooling ($r = 5$) | Attention pooling ($r = 5$) |

##### S.2.2.2 Gated Fusion Module

Let $\mathbf{H}_s, \mathbf{H}_x \in \mathbb{R}^{B\times d_h}$ denote the site-centric representations produced by the sequence and structure towers, respectively, and let $\mathbf{C}_s, \mathbf{C}_x \in \mathbb{R}^{B\times L\times d_h}$ denote the corresponding contextual feature maps.

The two modalities are first aligned through cross-modal attention. Taking the sequence branch as query and the structural branch as context, the refined sequence representation is obtained as

$$\tilde{\mathbf{H}}_s = \mathrm{LN}\big(\mathbf{H}_s + \mathrm{Attn}(\mathbf{H}_s W_Q^s,\ \mathbf{C}_x W_K^x,\ \mathbf{C}_x W_V^x)\big), \tag{S5}$$

and $\tilde{\mathbf{H}}_x$ is computed analogously with the roles of the two branches exchanged.

Because predicted structures may contain regions of low reliability, we introduce a confidence-based adaptive gating mechanism to modulate the structural signal before fusion. Concretely, a two-layer perceptron followed by learnable calibration produces a per-element confidence mask:

$$s_{\mathrm{raw}} = \sigma\big(W_2\, \mathrm{ReLU}(W_1 \tilde{\mathbf{H}}_x + \mathbf{b}_1) + b_2\big), \tag{S6}$$

$$\mathrm{conf} = \sigma(\alpha\, s_{\mathrm{raw}} + \beta), \qquad \hat{\mathbf{H}}_x = \tilde{\mathbf{H}}_x \odot \mathrm{conf}, \tag{S7}$$

where $\alpha$ and $\beta$ are learnable scalar parameters. This design allows the model to downweight unreliable structural predictions without discarding them entirely.

The modulated representations are then combined via a gated fusion layer. A gating vector $\mathbf{g}$ is predicted from the concatenation of the two branches, and the fused representation is computed as

$$\mathbf{g} = \sigma\big(\mathrm{MLP}_{\mathrm{gate}}([\tilde{\mathbf{H}}_s;\ \hat{\mathbf{H}}_x])\big), \quad \mathbf{Z}_{\mathrm{fused}} = W_{\mathrm{out}}\big[\mathbf{g} \odot \tilde{\mathbf{H}}_s;\ (1-\mathbf{g}) \odot \hat{\mathbf{H}}_x\big]. \tag{S8}$$

The fused representation $\mathbf{Z}_{\mathrm{fused}}$ is passed to a classification head for binary PTM-site prediction and an auxiliary regression head for modification-confidence estimation. The model is trained end-to-end by minimising a composite loss

$$\mathcal{L}_{\mathrm{total}} = \lambda\, \mathcal{L}_{\mathrm{focal}} + (1-\lambda)\, \mathcal{L}_{\mathrm{mse}}, \qquad \mathcal{L}_{\mathrm{focal}}(p_t) = -\alpha_t (1-p_t)^{\gamma} \log p_t, \tag{S9}$$

where $\alpha_t = 0.6$, $\gamma = 2.0$, and $\lambda = 0.7$. The focal loss addresses the severe class imbalance inherent in PTM-site prediction, while the mean-squared-error term regularises the confidence estimates produced by the gating mechanism.

##### S.2.3 PTM-Aware Dynamics Through Adaptive Conditioning

The PTM-Aware generator extends the base dynamics generator (Section S.2.1) with three targeted architectural modifications that collectively enable PTM-aware conformational ensemble generation while preserving the general dynamics priors acquired during pre-training. The core challenge is to integrate site-specific information into a pre-trained generative model without catastrophic forgetting or excessive computational overhead. Our solution follows a *minimal-perturbation* principle: the pre-trained IPA backbone, temporal attention layers, and flow-matching objective are retained unchanged; only a small number of new or expanded parameters are introduced, all carefully initialised to ensure functional equivalence with the pre-trained model at fine-tuning onset.

###### S.2.3.1 Modified-Residue Vocabulary Expansion

The base dynamics generator encodes each residue with a standard 21-class amino-acid vocabulary (20 canonical types plus an unknown token X). To natively represent phosphorylated residues, the embedding matrix is expanded from $\mathbb{R}^{21\times d}$ to $\mathbb{R}^{24\times d}$ by appending three phosphorylation types, phosphoserine (SEP, index 21), phosphothreonine (TPO, index 22), and phosphotyrosine (PTR, index 23). A dedicated token for each modification type allows the embedding layer to reflect the physicochemical consequences of phosphorylation within the learned representation space rather than relying solely on external conditioning signals.

New embedding vectors are initialised from their unmodified precursors (Ser→SEP, Thr→TPO, Tyr→PTR), so the initial representation of, e.g., phosphoserine is identical to serine and the model begins fine-tuning from a chemically meaningful starting point, needing only to learn the perturbation induced by the phosphate group. Compared with random initialisation, which would require simultaneously learning both canonical residue semantics and the modification effect, this precursor-based strategy leads to faster convergence. The EMA shadow-weight repair procedure that ensures consistency of the expanded embedding matrix with the exponential moving average is detailed in Section S.3.3.

###### S.2.3.2 PTM Adapter

A central design question is how to inject PTM-specific information into a pre-trained dynamics generator without disrupting the general protein dynamics priors it has already acquired. Direct modification of the backbone weights risks catastrophic forgetting; conversely, a purely external conditioning signal may be too weak to capture the subtle structural perturbations induced by phosphorylation. We adopt a lightweight adapter strategy that balances these concerns.

Each residue $i$ receives a 256-dimensional PTM feature vector $\mathbf{f}_i \in \mathbb{R}^{256}$ extracted from the gated fusion layer of the PTM prediction module (Section S.2.2). Importantly, *every* residue, whether modified or not, passes through the same feature extraction pipeline; no manual zeroing or binary masking is applied. The resulting features encode rich, site-specific information including sequence context motifs, local structural geometry, and cross-modal confidence scores, all of which are distilled during PTM-site prediction training. Because the PTM prediction module is trained to discriminate modified from unmodified sites, the learned representations naturally embed this distinction: features at modification sites carry PTM-specific activation patterns, while features at non-modification sites reflect the local sequence and structure context without such patterns. This implicit differentiation enables the downstream dynamics generator to selectively respond to PTM signals without requiring an explicit binary mask or auxiliary masking loss.

The feature vector is projected to the backbone hidden dimension $d$ via a two-layer MLP adapter with SiLU activation:

$$\mathbf{e}_i^{\text{PTM}} = W_2\,\text{SiLU}(W_1\mathbf{f}_i + \mathbf{b}_1) + \mathbf{b}_2, \quad W_1 \in \mathbb{R}^{d\times 256},\ W_2 \in \mathbb{R}^{d\times d}. \tag{S10}$$

Critically, the output projection $(W_2, \mathbf{b}_2)$ is *zero-initialised*: at fine-tuning onset the adapter output is identically zero for *all* residues (modified or not), so the model's forward pass is mathematically identical to the pre-trained checkpoint. As gradients flow through the adapter during fine-tuning, the PTM signal is progressively activated—a "warm-start" property that prevents training instability and ensures that the pre-trained dynamics quality is preserved in early training iterations.

###### S.2.3.3 Conditioning Injection

PTM-induced conformational changes span multiple spatial and temporal scales: local effects include altered backbone dihedral preferences and side-chain reorientation at the modification site, while long-range (allosteric) effects can shift domain–domain interfaces or modulate loop flexibility tens of Å away. A single injection point is unlikely to capture both regimes simultaneously. We therefore introduce a two-level conditioning scheme that injects PTM information at complementary stages of the forward pass.

**Sequence embedding fusion.** The adapter output is added residually to the amino-acid embedding before the IPA stack:

$$\tilde{\mathbf{h}}_i^{\text{aa}} = \text{Emb}_{\text{aa}}(s_i) + \mathbf{e}_i^{\text{PTM}}. \tag{S11}$$

Because IPA layers propagate information through geometric message-passing, a perturbation at the embedding level is amplified through successive layers and can influence distant residues via the spatial attention mechanism. This level primarily modulates the *per-residue identity* seen by the backbone. The model effectively perceives a phosphoserine as a distinct residue type whose embedding differs from serine by the learned adapter offset, enabling it to predict site-local conformational shifts.

**Temporal latent injection.** After IPA processing, the spatially contextualised output $\mathbf{x}^{\text{IPA}} \in \mathbb{R}^{B \times L \times d}$ already encodes the structural consequences of PTM conditioning propagated through the geometric message-passing layers. This representation is injected into the temporal latent tensor via residual addition, broadcast across all $T$ trajectory frames:

$$X^{\text{main}} \leftarrow X^{\text{main}} + \mathbf{x}^{\text{IPA}}[\,\cdot, \text{None}, \cdot, \cdot\,]. \tag{S12}$$

The broadcasting operation ensures that the PTM-conditioned spatial context is shared uniformly across the temporal dimension, providing a consistent structural prior for all generated frames. The temporal attention layers can learn how PTM-induced structural perturbations translate into altered fluctuation amplitudes, modified correlated motions, and shifted conformational equilibria across the entire trajectory.

###### S.2.3.4 PTM-Conditioned Inference Pipeline

At inference time, the PTM-conditioned generator produces an all-atom conformational ensemble in four stages. The pipeline is designed to be modular: the PTM prediction module runs once to extract per-residue features, which are then consumed by the dynamics generator without further gradient computation.

**Input preparation.** The input sequence is modified by replacing canonical residue tokens at annotated PTM sites with the corresponding expanded-vocabulary tokens (SEP/TPO/PTR). All residues are then processed through the frozen PTM prediction module's gated fusion layer to obtain 256-dim feature vectors $\{\mathbf{f}_i\}$, which are projected via the adapter (Eq. S10). Consistent with the training pipeline, every residue receives a non-zero feature vector encoding its local sequence and structural context; the distinction between modified and unmodified sites is implicitly captured by the learned activation patterns of the PTM prediction module rather than by explicit masking.

**Structure conditioning.** Atom14 heavy-atom coordinates and per-residue rigid-body frames are extracted from the reference structure $T^{(0)}$. Backbone torsion angles $(\phi, \psi, \omega)$ and side-chain dihedrals $(\chi_1$–$\chi_4)$ are computed from the atom14 representation and encoded as sine–cosine pairs. These structural features are combined with the PTM embeddings from Step 1 through the two-level injection scheme (Eqs. S11–S12).

**ODE integration.** Starting from Gaussian noise $\mathbf{x}_0 \sim \mathcal{N}(\mathbf{0}, \mathbf{I})$, Eq. S4 is integrated under PTM conditioning using the Dormand–Prince adaptive solver (`dopri5`) with tolerances $\varepsilon_{\text{abs}} = 10^{-6}$, $\varepsilon_{\text{rel}} =$

$10^{-3}$. The PTM conditioning enters the velocity field $v_\theta$ at every solver evaluation through both injection levels, ensuring that the modification signal consistently guides trajectory generation throughout the integration path.

**Structure reconstruction.** The generated latent trajectory $\mathbf{x}_1$ is decoded to all-atom coordinates via quaternion-to-rotation conversion and frame composition (Algorithm S2). Before atom14 decoding, modified tokens are mapped back to their canonical residue types (SEP→Ser, TPO→Thr, PTR→Tyr) to ensure correct heavy-atom coordinate lookup from the standard residue geometry library. This reverse-mapping is essential because the atom14 scheme indexes atoms by canonical residue type; without it, the decoder would attempt to look up atom positions for non-existent residue types, producing coordinate errors.

#### S.3 Training Methodology

##### S.3.1 Pre-training: Multi-scale Strategy and Hyperparameters

Protein conformational dynamics span a broad hierarchy of timescales. Local motions such as side-chain rotamer flips and loop fluctuations occur on the picosecond-to-nanosecond range, whereas domain hinge movements and large-scale folding–unfolding transitions take place over tens to hundreds of nanoseconds or longer. A single temporal resolution cannot faithfully capture both regimes simultaneously, so we pre-train the base dynamics generator at two complementary resolutions in independent runs that share an identical model architecture, the same hyperparameters, and the same sampling procedure. In the short-timescale (100 ps) stage, ATLAS and MoDEL trajectories are sub-sampled at 100 ps intervals so that each training sample comprises 100 consecutive frames spanning a 10 ns observation window; this resolution is also adopted during PTM-conditioned fine-tuning so as to match the DynaMo-phos dataset, thereby eliminating any resolution gap at transfer time. In the long-timescale (1 ns) stage, ATLAS and mdCATH trajectories are sub-sampled at 1 ns intervals, yielding 100-frame windows that cover 100 ns and are therefore better suited to capturing slow, large-amplitude collective motions; this stage also provides a foundation for future applications that require modelling of slower dynamical processes. Training the two stages independently allows each model to focus on a single characteristic timescale, avoiding gradient conflicts that would arise from mixing samples whose inter-frame displacements differ by an order of magnitude, and thus improving learning efficiency.

To prevent any single large dataset from dominating the training distribution, we employ a hierarchical stratified sampling strategy. At each iteration a source dataset is first drawn according to fixed weights

(ATLAS : MoDEL = 0.5 : 0.5 for the 100 ps stage; ATLAS : mdCATH = 0.5 : 0.5 for the 1 ns stage), after which a protein system is selected uniformly at random from the chosen dataset and a start frame is drawn uniformly along the trajectory. A contiguous block of 100 frames beginning at the selected position is then extracted and featurised; systems whose trajectories contain fewer than 100 frames are excluded automatically. The complete set of pre-training hyperparameters is listed in Table S5.

**Table S5: Pre-training hyperparameters.** Architecture and optimisation settings are shared across both temporal stages; only data composition differs.

| Category | Hyperparameter | Value |
|---|---|---|
| **Architecture** | Hidden dimension $d$ | 384 |
| | Transformer layers | 5 |
| | MHA heads | 16 |
| | IPA heads | 4 |
| | IPA head dim. | 32 |
| | IPA query/key dim. | 8 |
| | IPA value dim. | 8 |
| **Optimisation** | Optimiser | AdamW |
| | Learning rate | $1 \times 10^{-4}$ |
| | EMA decay | 0.999 |
| | Gradient clipping (max norm) | 1.0 |
| | Gradient accumulation | 1 |
| | Precision | float32 |
| **Schedule** | Epochs | 1000 |
| | Batch size (per GPU) | 1 |
| **Data** | Frames per sample | 100 |
| | Max residues (crop) | 256 |
| | 100 ps datasets | ATLAS + MoDEL (0.5 : 0.5) |
| | 1 ns datasets | ATLAS + mdCATH (0.5 : 0.5) |
| **Flow matching** | Path type | GVP |
| | Prediction target | velocity |
| | Time multiplier | 100 |
| | $\alpha_{\max}$ (torsion) | 8 |
| | Discrete loss weight | 0.5 |
| **Inference (ODE)** | Solver | `dopri5` |
| | $\varepsilon_{\text{abs}}$ | $1 \times 10^{-6}$ |
| | $\varepsilon_{\text{rel}}$ | $1 \times 10^{-3}$ |
| **Hardware** | GPU | NVIDIA A100 40 GB |

##### S.3.2 PTM Site Prediction

All PTM prediction experiments were conducted on NVIDIA RTX 3090 (24 GB) GPUs using Python 3.10, PyTorch 2.1, CUDA 11.8, and fair-esm 2.0.0, consuming approximately 1,200 GPU-hours in total across all 18 PTM types. The full set of hyperparameters is provided in Table S6.

**Table S6: PTM site prediction hyperparameters.**

| Category | Hyperparameter | Value |
|---|---|---|
| **Architecture** | Hidden dimension $d_h$ | 256 |
| | Transformer layers (Seq/Struct) | 4 |
| | Attention heads | 16 |
| | MoE experts | 6 |
| | ESM-2 feature dim. | 1280 |
| | ESM-IF1 feature dim. | 512 |
| | Dropout | 0.25 |
| **Optimisation** | Optimiser | AdamW |
| | Learning rate | $3 \times 10^{-5}$ |
| | Weight decay | 0.1 |
| | Scheduler | Cosine with warmup |
| | Warmup steps | 500 |
| | Gradient accumulation | 2 |
| | Precision | float32 |
| **Schedule** | Epochs | 50 |
| | Batch size | 256 |
| | Early stopping patience | 5 |
| **Data** | Window size | 73 |
| | Local window size | 21 |
| | Centre window radius | 5 |
| | Augmentation probability | 0.5 |
| | Feature noise level | 0.15 |
| **Loss** | Focal loss $\alpha$ | 0.6 |
| | Focal loss $\gamma$ | 2.0 |
| **Hardware** | GPU | NVIDIA RTX 3090 24 GB |

##### S.3.3 PTM-Conditioned Fine-Tuning

Fine-tuning is initialised from the 100 ps-resolution pre-trained model. We employ a full-parameter fine-tuning approach, training all model components end-to-end without layer freezing. This includes the Invariant Point Attention (IPA) backbone, the temporal attention mechanism, the expanded embedding matrix, and the newly introduced PTM adapter. Pre-trained weights are loaded using a shape-aware strategy that accommodates naming remapping and elegantly handles shape-mismatched layers, most notably the expanded embedding matrix (from $21 \times d$ to $24 \times d$). To ensure stable training and preserve the general dynamics priors learned during pre-training, the PTM adapter parameters are initialized with zeros, enforcing a gradual-activation property. We use a reduced learning rate of $5 \times 10^{-5}$ for all parameters, which allows the adapter to converge rapidly while mitigating catastrophic forgetting of the established representation.

Expansion of the residue embedding matrix prior to fine-tuning introduces a shape mismatch with the EMA ($\gamma = 0.999$) shadow weights. To resolve this, a two-stage repair procedure is applied immediately after vocabulary expansion. First, the entire model state is hard-copied into the EMA shadow buffer, thereby eliminating all dimensional inconsistencies in a single global synchronisation step. The shadow embeddings corresponding to the three newly introduced PTM tokens are then overwritten with the EMA embeddings of their respective precursor residues (Ser, Thr and Tyr), ensuring that the shadow representation preserves the learned semantic continuity of the unmodified amino acids. All inference is performed exclusively with EMA shadow weights.

#### S.4 Conformational Dynamics Benchmarking

This section systematically evaluates the flow-matching backbone on established molecular-dynamics benchmarks, examining the effects of temporal resolution, architectural components and input representations on the fidelity of generated conformational ensembles.

##### S.4.1 Evaluation Protocol

All metrics are computed on C$\alpha$ coordinates after superposition onto the reference crystal structure. We denote by $T$ the number of frames, $L$ the number of residues, and $\mathbf{r}_i(t)$ the C$\alpha$ position of residue $i$ at frame $t$.

**Structural diversity. Pairwise RMSD correlation.** For each protein in the test set, we compute the mean pairwise C$\alpha$ RMSD across all frame pairs within the generated ensemble and within the reference MD ensemble. The Pearson correlation of these per-protein values across the test set measures whether the model reproduces the rank ordering of conformational heterogeneity.

**Local flexibility.** **Per-target RMSF Pearson $r$.** Per-residue root-mean-square fluctuation around the ensemble mean is defined as

$$\mathrm{RMSF}_i = \sqrt{\frac{1}{T}\sum_{t=1}^{T}\left\|\mathbf{r}_i(t) - \langle \mathbf{r}_i \rangle\right\|^2}. \tag{S13}$$

We report the median of per-protein Pearson correlations between generated and reference RMSF profiles across the test set. For the BK channel case study, we additionally report RMSE between the two RMSF profiles as a measure of absolute error.

**RMSD coverage.** For each protein in the test set, we compute the fraction of reference MD frames for which at least one generated frame falls within a C$\alpha$ RMSD threshold of 1.0 Å after superposition:

$$\text{Coverage} = \frac{1}{T}\sum_{t=1}^{T}\mathbf{1}\left[\min_{s}\ \mathrm{RMSD}(\mathbf{r}(t),\ \hat{\mathbf{r}}(s)) < 1.0\,\text{Å}\right]. \tag{S14}$$

We report the mean coverage across the test set.

**Conformational landscape.** Free energy landscapes are estimated by projecting generated and reference MD ensembles onto the first two principal components (PCs) of the reference MD trajectory, computing the negative log-probability density via kernel density estimation, and reporting values in kcal/mol. Kinetic properties are assessed via time-lagged independent component analysis (tICA) with a lag time of 10 ns, projecting onto the two slowest independent components to visualise conformational transition pathways.

#### S.5 PTM Site Prediction Benchmarking

This section reports the classification performance of the dual-modal PTM site prediction module across 18 post-translational modification types, together with comparisons against published structure-aware baselines and internal ablation analyses.

##### S.5.1 Evaluation Metrics

Classification performance is quantified by accuracy (ACC), sensitivity (SN), specificity (SP) and Matthews correlation coefficient (MCC), defined as:

$$\mathrm{ACC} = \frac{TP + TN}{TP + TN + FP + FN}, \tag{S15}$$

$$\mathrm{SN} = \frac{TP}{TP + FN}, \tag{S16}$$

$$\mathrm{SP} = \frac{TN}{TN + FP}, \tag{S17}$$

$$\mathrm{MCC} = \frac{TP \cdot TN - FP \cdot FN}{\sqrt{(TP + FP)(TP + FN)(TN + FP)(TN + FN)}}, \tag{S18}$$

where $TP$, $TN$, $FP$, and $FN$ denote true positives, true negatives, false positives, and false negatives, respectively. Statistical significance is assessed via non-parametric bootstrap resampling ($n$ = 1,000 iterations; 95% confidence intervals by the percentile method) and the Wilcoxon signed-rank test ($P < 0.05$).
